# Comparison of the performance of sporophytes derived from narrow and wide genetic crosses between strains of the cultivated kelp *Saccharina latissima*

**DOI:** 10.64898/2026.10.12.758020

**Authors:** Zofia Nehr, Francis Le Madec, Olivier Godfroy, Aurélie Préchoux, Alan Le Moan, Komlan Avia, Dan Potin, Martina Strittmatter, Marie-Mathilde Perrineau, Cecilia Rad-Menendez, Delphine Scornet, Bertrand Jacquemin, Emilie Gouhier, Jerome Coudret, Lucie Jaugeon, Stephane Mauger, Carla Ruiz Gonzalez, Myriam Valero, Claire Gachon, Philippe Potin, J. Mark Cock

## Abstract

Seaweed mariculture allows sustainable production of large quantities of biomass for multiple downstream applications but the genotypes cultivated by this industry need to be improved to increase efficiency and cost-effectiveness. It is important that the genetic improvement of cultivated strains does not lead to the introduction of foreign genetic material into wild seaweed populations, which are already under threat from global heating. To circumvent this problem, efforts are being made to develop breeding approaches based on the use of locally-resourced genetic material. The aim of this study was to evaluate the potential effectiveness of this ‘local’ breeding strategy compared to a classical breeding approach based on wide genetic crosses. Analyses of growth rates and cell wall composition of F1 hybrids derived from either intra- or inter-population crosses between *Saccharina latissima* strains from two localities, southern Brittany, France and the Faroe Islands, did not identify any clear advantage of inter-compared to intra-population crosses. Breeding based on locally-resourced strains would therefore appear to be a valid approach for current strain selection programs. The study also established a large collection of gametophyte strains from multiple sites for future breeding programs and showed that these strains retained the capacity to become fertile after more than ten years in culture.

## INTRODUCTION

Seaweed mariculture is a highly sustainable approach to the production of plant biomass for food, feed and other downstream applications (Buschmann *et al*., 2017; Cai *et al*., 2021; Sæther *et al*., 2024; United Nations, 2024). Seaweed cultivation does not require freshwater nor arable land and rapid growth rates mean that these organisms can potentially produce very large quantities of biomass per hectare. These characteristics have led to the development of large-scale seaweed farms in Asia, where production by cultivation greatly exceeds the biomass available from harvesting wild algae (Verdegem *et al*., 2023). In contrast, in Europe and northern America where the most commonly cultivated species is the sugar kelp *Saccharina latissima* (Linnaeus) C.E. Lane, C. Mayes, Druehl & G.W. Saunders (Lane *et al*., 2007), seaweed mariculture is still an emerging sector and the majority of macroalgal biomass is obtained by harvesting wild populations (Sæther *et al*., 2024). Future upscaling of the seaweed mariculture sector in the western hemisphere will require significant research and development investment at multiple levels (production, harvesting, processing, marketing, etc.) but improvement of yield and biomass quality, as a starting point for the value chain, is a particularly important target. One promising approach in this context is the application of breeding approaches to develop strains with improved yields or other characteristics of interest such as biomolecule content or stress tolerance. There is considerable scope for improvement of strains through breeding in Europe and northern America because seaweed farms principally use wild-sourced strains that have not been subjected to any genetic improvement through breeding. Although there is a long history of breeding being applied to the closely related kelp *Saccharina japonica* in Asia (Fang and Li, 1966; Tseng, 2001; Li *et al*., 2007, 2008, 2016; Zhang *et al*., 2007, 2011; Liu *et al*., 2010, 2014; Hwang *et al*., 2019; Coelho and Cock, 2020; Hu *et al*., 2024), initiatives aimed at breeding improved strains of *S. latissima* in Europe and northern America are much more recent (Goecke *et al*., 2020; Umanzor *et al*., 2021; Huang *et al*., 2022, 2023; Li *et al*., 2022; Cohen *et al*., 2025; Gonzalez *et al*., 2025; Nehr *et al*., 2025; Bråtelund *et al*., 2026b, 2026a; Mauger *et al*., 2026).

It is important that upscaling of seaweed mariculture does not lead to negative impacts on the marine environment. One current concern is that the development of large-scale seaweed farms based on genetically-improved strains may have a negative impact on wild populations through the introduction of foreign genetic material via escapes from farms (Campbell *et al*., 2019). This is an important consideration, given the current fragile state of wild seaweed populations due to the effects of global heating (Wernberg *et al*., 2011; Smale, 2020). To avoid the introduction of non-local genotypes, selection programs would need to use only locally-sourced genetic material to produce improved strains to be grown in a specific region. Indeed, the regulatory frameworks that govern seaweed cultivation in north Atlantic most countries typically restrict the use of non-locally-sourced or non-native strains in order to prevent the introduction of non-indigenous genotypes into coastal ecosystems (Goecke *et al*., 2020). Consequently, breeding programs cannot rely on importing elite strains from other regions, a practice common in terrestrial crop improvement, but must instead exploit genetic diversity present within local or regional wild populations. This makes the systematic sampling, genotyping, and preservation of wild *S. latissima* germplasm across its native range a critical first step (Hofmann *et al*., 2025), both to assess the breadth of available diversity and to constitute living collections of geographically referenced, characterised material that breeders can legally and practically access. Several studies have investigated the genetic diversity and structure of *S. latissima* across its range using microsatellite markers and, more recently, genome-wide SNP approaches (Breton *et al*., 2018; Luttikhuizen *et al*., 2018; Mooney *et al*., 2018; Neiva *et al*., 2018; Guzinski *et al*., 2020; Mao *et al*., 2020; Grant and Chenoweth, 2021; Diehl *et al*., 2023; Jaugeon *et al*., 2025; Puncher *et al*., 2026; Treillefort *et al*., 2026). These studies have consistently identified significant genetic differentiation between northern and southern European populations, a latitudinal gradient in within-population diversity, and evidence of local adaptation to environmental variables including sea surface temperature and salinity. It is important that biobanking of genetic resources for *S. latissima* takes into account its Europe-wide genetic structure to ensure that appropriate material is available for future breeding programs.

The objective of this study was to evaluate to what extent the use of locally-sourced progenitor genotypes might limit (or not) the efficacity of future breeding programs, compared to more classical approaches based on wide crosses (Cohen *et al*., 2025). For this, we first established a large, diverse collection of 650 *S. latissima* gametophyte strains. Two local sites were then selected - the southern part of the Finistère department of France and the Faroe Islands - and both intra-population and inter-population crosses were carried out using gametophyte strains from the two sites. Comparative analysis of the growth and cell wall characteristics of the resulting F1 hybrid sporophytes did not identify any significant advantage of wide crosses over crosses between locally-sourced strains in these experiments supporting current initiatives aimed at promoting the use of the latter in seaweed breeding programs.

## MATERIALS AND METHODS

### Sample collection and isolation of gametophyte strains

Isolated female and male gametophyte strains were derived from fertile *S. latissima* blades collected in the field at 33 different sites across Europe over the course of two sampling campaigns carried out in the context of projects financed by the European Union, the first from 2014 to 2019 for the project Genialg (grant agreement number 727892) and the second from 2022 to 2023 for the project SeaMark (grant agreement number 101060379).

During the first campaign, fertile tissue was collected by excising approximately 1 cm² explants from the fertile region of the blade (sorus). Explants were briefly rinsed with fresh water and gently wiped with absorbent paper to remove surface epiphytes. Two explants per individual were placed on either side of two glass slides within a 50 mL Falcon tube filled with 45 mL of filter-sterilised seawater. Tubes were transported to the laboratory in insulated boxes with ice packs to maintain a low temperature during transit. Upon arrival, tubes were incubated in the dark at 13°C under gentle agitation (approximately 150 rpm) for 6 to 8 hours to allow spore release. Following spore release, the suspension from each sporophyte was distributed into three 90 mm Petri dishes. The two glass slides were each transferred to a separate dish, and all three dishes were topped up to volume with half-strength Provasoli-enriched (Starr and Zeikus, 1993) natural seawater (PES).

During the second campaign, portions of fertile blades were transported to the laboratory in insulated boxes with ice packs. Upon arrival, epiphytes were removed from the blade surface using a razor blade, and the blade tissue was wiped with absorbent paper. Small pieces of approximately 1 mm² were then dissected from the fertile region and rinsed with sterile seawater. Each piece was placed in a 40 µL drop of sterile seawater on a glass coverslip positioned at the bottom of a 55 mm Petri dish. The dish was inverted and exposed to white light for at least 3 hours to induce spore release onto the coverslip, (adapted from Coelho *et al*., 2012). Once spores had settled and attached to the coverslip, it was carefully removed, gently rinsed in sterile seawater, and transferred to a clean 55 mm Petri dish containing half-strength PES prepared with natural sterile seawater.

For both sampling campaigns, gametophyte pools were then maintained for one month under red light with a 16h:8h light:dark photoperiod at 13°C before isolation. Red light illumination was from Osram 18W/840 fluorescent “day light” neon tubes (Osram GmbH, Germany) filtered through primary red lighting gelatine sheet (reference 106, Lee Filters, United Kingdom). Individual gametophytes were then detached from the surface of the Petri dish, pipetted in 20 µL of medium and transferred to a 24-well cell culture plate where they were supplemented with additional PES and grown under the same red light conditions. This procedure established a collection of 650 *S. latissima* gametophyte strains from 33 European sites (Fig. 1).

**Fig. 1.**
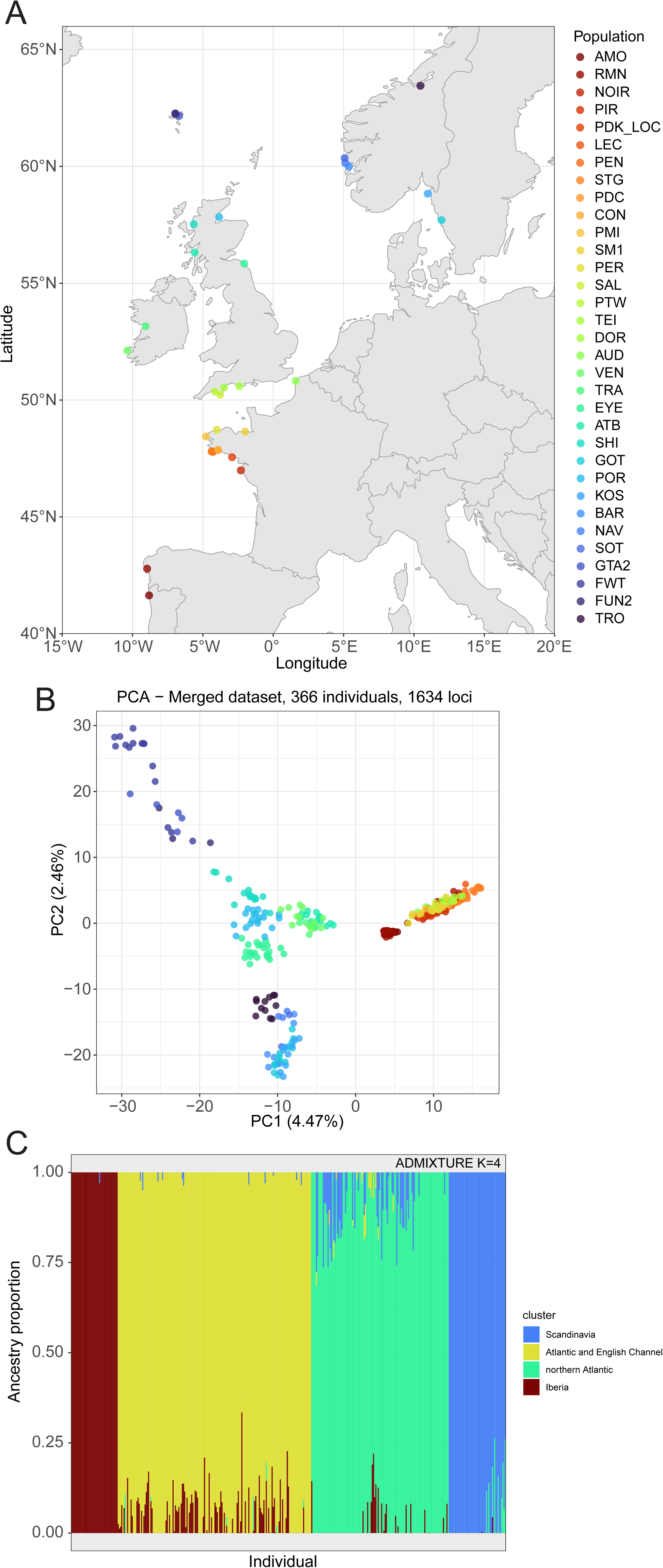
Sampling and genetic analysis of a *S. latissima* gametophyte collection. A. Locations of the 33 sampling sites (see Table S4 for details). B. Principal component analysis of 366 individuals from the sampled populations based on 1,634 SNPs. Population colour codes are the same as for panel A. The first two principal components explain 4.47% and 2.46% of the total variance, respectively. C. Population structure based on ADMIXTURE analysis. Each vertical bar represents an individual gametophyte and shading indicates proportional membership to the four colour-coded clusters. AMO, Amorosa; RMN, Ría de Muros y Noya; NOIR, Pointe Devin, Noirmoutier; PIR, Piriac-sur-Mer; PDK_LOC, Locmariaquer; LEC, Lechiagat; PEN, Penmarc’h; STG, Saint-Guénolé; PDC, Point de Cabellou; CON, Concarneau; PMI, Lanildut; SM1, Saint Malo; PER, Perharidy; SAL, Salcombe; PTW, Plymouth; TEI, Teignmouth; DOR, Dorset, Portland Harbour; AUD, Audresselles; VEN, Ventry Bay; TRA, Ballyvelaghan; EYE, Eyemouth; ATB, Atlantic Bridge (Clach Na Sula); SHI, Shieldaig, Strathcarron; GOT, Göteborg; POR, Portmahomack, Tain; KOS, Kosterhavet National Park; BAR, Bårdsundet; NAV, Naveide; SOT, Sotra area; GTA2, rig AA03, Gøtuvík; FWT, Funningsfjørður; FUN2, Funningsfjørður A02; TRO, Ringvebukta, Trondheim.

### PCR-based determination of gametophyte sex

After about six months in culture, a small piece of gametophyte thallus was removed, cut into small pieces under a binocular microscope and then transferred to a 1.5 mL tube where the medium was adjusted to a final volume of 100 µL with milliQ water. The samples were frozen and thawed twice before carrying out the PCR. PCR reaction mixes were 1x GoTaq buffer, 0.26 mM MgCl_2_, 0.04 mM for each of the four dNTPs, 0.02 mg/ml BSA, 0.12 mM of each PCR primer (Table S1) and 0.03 units/µL of GoTaq® G2 Flexi DNA polymerase (Promega) in a final volume of 12.5 mL and the PCR amplification protocol was as follows: 96°C for 3 min, followed by 10 cycles of 96°C for 40 sec, 65°C (-1°C/cycle) for 40 sec and 72°C for 40 sec, then 25 cycles of 96°C for 40 sec, 55°C for 40 sec and 72°C for 40 sec, and finally 72°C for 10 min. A sample of 12.5 µL was then analysed by electrophoresis on a 2.5% agarose gel with appropriate size markers. Each PCR reaction contained a pair of male-specific primers, a pair of female-specific primers and a pair of positive control primers which amplified the mitochondrial cytochrome c oxidase I (coxI-5′) gene (Lane *et al*., 2007). The sex-specific primers correspond to markers that were developed originally to detect sex-linked scaffolds in the closely related species *Saccharina japonica* (Table S1; Lipinska *et al*., 2017).

### Genotyping of gametophyte strains

Gametophytes were genotyped using the ddRAD-seq method (Peterson *et al*., 2012) applying the protocol described by Mauger and Avia (2024). Genotyping involved the construction and sequencing of two ddRAD-seq library pools (ddRAD pool 1 and ddRAD pool 2) at different time periods. ddRAD pool 1 included 409 samples from the Genialg campaign, whereas ddRAD pool 2 included 138 samples from the Seamark campaign plus 36 samples from the Genialg campaign that had not been genotyped previously. In total, 583 gametophyte strains were genotyped using ddRAD-seq (all 444 strains from the Genialg campaign and 139 of the 206 strains from the SeaMark campaign).

For DNA extraction, a mass of around 1 mm² of gametophyte filaments was harvested, blotted dry on absorbent paper, placed in a 1.5 mL microtube and immediately frozen in liquid nitrogen. The tissue was disrupted with four pulses of 45 sec in a Qiagen TissuLyzer II (Qiagen, Hilden, Germany; ddRAD pool 1 samples) or manually disaggregated in a microtube using a micropestle (ddRAD pool 2 samples). DNA was extracted using the NucleoSpin Plant II 96 kit (Macherey-Nagel GmbH & Co. KG, Düren, Germany). Lysis was carried out in PL1 buffer at room temperature for 15 to 30 min with vigorous agitation to resuspend all the disrupted tissue. Additional washes with PW1 (in some cases) and PW2 buffer were carried out at the end of the extraction to further improve elimination of PCR inhibitors. DNA was eluted into 60 µL or 100 µL of PE buffer at 70°C. ddRAD pool 1 samples were further purified and concentrated using the NucleoSpin® gDNA Clean-up XS micro kit (Macherey-Nagel GmbH & Co. KG, Germany) according to the manufacturer’s instructions, with elution into 30 μL (2 x 15 µL) of the supplied elution buffer. DNA concentration was measured using PicoGreen (Fisher Scientific) and DNA integrity was evaluated on a 1.5% agarose gel. For samples purified with the NucleoSpin® gDNA Clean-up XS micro kit, the concentration of the purified DNA was adjusted to 2.5 ng.µL^-1^.

ddRAD pool 1 genomic DNA samples were digested with *Pst*I and *Hha*I and ddRAD pool 2 genomic DNA samples were digested with *Pst*I and *Mse*I (New England Biolabs, https://www.neb.com). Samples of 100 ng of DNA from individual gametophyte strains were digested overnight at 37°C with 10 U of each enzyme in 1X CutSmart buffer (total volume of 50 µL). Digested DNA was purified with AMPure XP beads (Beckman Coulter Genomics). Adapters were added using T4 DNA ligase (New England Biolabs). Illumina flowcell annealing sequences, multiplexing indices and sequencing primer annealing regions were added to all fragments bearing adapters by PCR amplification using the Q5 Hot Start High-Fidelity DNA polymerase kit (New England Biolabs) so that each individual library could be identified with a unique combination of barcode and index. Sixteen barcodes and either 24 (ddRAD pool 1) or 12 (ddRAD pool 2) Illumina indexes were used. The concentration of each library was checked on a 1.5% agarose gel. For ddRAD pool 1, individual libraries were pooled into three intermediate concentration groups (low, medium, and high), concentrated with AMPure XP beads (Beckman Coulter Genomics), and quantified using the Qubit dsDNA HS Assay kit (Thermo Fisher Scientific) to enable equimolar pooling. For ddRAD pool 2, 5 µL aliquots of each individual library were pooled and precipitated with 2.5 volumes of ethanol and 0.1 volume of 3 M sodium acetate (pH 5.2). Fragments of between 480 and 930 bp (ddRAD pool 1) or between 480 and 980 bp (ddRAD pool 2) were then selected with a Pippin-Prep kit (Sage Science, Beverly, MA, USA). An Agilent 2100 Bioanalyzer (Agilent Technologies) was used to evaluate the insert size ranges of the final library pools and the library pools were quantified using the Qubit DNA HS kit (Invitrogen). The two final ddRAD pools were sequenced at Novogene (Munich, Germany) on an Illumina HiSeq 2500 chemistry V4 run (ddRAD pool 1) or on a partial lane of an Illumina NovaSeq X plus (ddRAD pool 2) to produce 125 bp (ddRAD pool 1) or 150 bp (ddRAD pool 2) paired-end reads.

Data were demultiplexed using the process_radtags module of Stacks version 2.65 (Catchen *et al*., 2011, 2013). Read quality was assessed and low-quality bases and adapter sequences were trimmed using fastp version 1.0.1 (Chen, 2025). Trimmed reads were mapped to the *S. latissima* chromosome-level v2 genome assembly (available at https://phaeoexplorer.sb-roscoff.fr/) using BWA version 0.7.17 (Li and Durbin, 2009). Mapped reads were sorted using SAMtools version 1.15.1 and SNPs were called using the ref_map.pl pipeline available in Stacks version 2.65 (Catchen *et al*., 2011, 2013). Only sequenced ddRAD fragments (RAD-tags) present in all the sampling sites, in at least 80% of the individuals and with a maximum heterozygosity of 80% (to remove potential paralogues) were retained for the downstream analysis, through the option of the Stacks module ‘populations’. The two resulting Variant Call Format (VCF) files, which initially identified 210,754 SNPs for the ddRAD pool 1 and 267,561 SNPs for ddRAD pool 2, were further filtered using VCFtools version 0.1.16 (Danecek *et al*., 2011) to retain only biallelic SNP loci with a minimum sequencing depth of five, a maximum sequencing depth of 40 and a minimum allele count of 2. The two filtered VCF files contained 141,735 SNPs for ddRAD pool 1 and 214,915 SNPs for ddRAD pool 2. Only a subset of the RAD-tags was shared between the datasets for ddRAD pool 1 and ddRAD pool 2 because the ddRAD sequencing employed the same infrequently cutting (6-base) restriction enzyme (*Pst*I) but used different frequently cutting (4-base) restriction enzymes (*Hha*I and *Mse*I for ddRAD pools 1 and 2, respectively). We therefore merged the two datasets based on common loci using the reference genome coordinates and the bcftools v1.16 isec function, resulting in a shared dataset of 1,634 SNPs. The project-specific datasets and the merged dataset were then analysed in parallel for population genetic analyses to assess the consistency of the results across datasets.

### Analysis of genetic diversity and genetic structure

Analyses of genetic diversity were carried out in R version 4.4.1 (R Core Team, 2024) using the packages vcfR version 1.16.0 (Knaus and Grünwald, 2017) for VCF file import, adegenet version 2.1.11 (Jombart and Ahmed, 2011) for genotype data manipulation, and ggplot2 version 4.0.3 (Wickham *et al*., 2016) for graphic visualisation. Prior to filtering, missing data and heterozygosity were quantified both per individual and per locus. This was carried out both for the gametophyte ddRAD-seq data and for a reference diploid sporophyte dataset (https://doi.org/10.5061/dryad.1jwstqjt8; Guzinski *et al*., 2020) as a diploid control. Missing data were computed as the proportion of loci with no genotype call per individual and as the proportion of individuals with no genotype call per locus. Observed heterozygosity was calculated per individual as the proportion of heterozygous loci, and per locus as the proportion of heterozygous individuals.

A high mean level of heterozygosity per individual was detected for the SeaMark dataset (1.91%), with particularly high values for the Norwegian populations BAR, NAV, SOT and the Faroe Islands population FUN2 (Table S2). As gametophytes are haploid organisms, detection of heterozygosity at a given locus can be an indication of contamination (i.e. a non-clonal gametophyte isolate) but may also be due to genotyping errors or PCR artefacts. These elevated levels of heterozygosity required the application of a stringent heterozygosity-based filtering step prior to downstream analyses.

Filtering was applied independently for the Genialg and SeaMark datasets in four sequential steps. First, individuals with more than 5% missing data were removed. Second, loci with more than 5% missing data (computed on the individual-filtered dataset) were removed. Third, individuals with more than 1% heterozygosity were removed. Fourth, loci with more than 1% heterozygosity were removed. To limit the effect of linkage disequilibrium between SNPs originating from the same RAD-tag, a single SNP was randomly retained per RAD-tag using a fixed seed (set.seed(42)) for both the SeaMark and Genialg datasets, after all filtering steps had been carried out. RAD-tag identity was inferred from the locus ID field of the VCF files, where the first numeric component corresponds to the catalogue locus identifier generated by Stacks version 2.65. The filtered datasets retained 363 individuals genotyped at 16,409 loci for ddRAD pool 1 and 141 individuals genotyped at 13,251 loci for ddRAD pool 2. The merged dataset was filtered using the same protocol, resulting in a filtered dataset of 366 individuals genotyped at 1,634 loci.

Principal Component Analysis (PCA) was carried out on mean-imputed, centred genotypes with the ade4 package version 1.7.24 (Dray and Dufour, 2007), retaining the first three principal components. Contributions of individual SNPs to the two first principal components of the PCA were calculated using extracted loadings. The contributions of each SNP to the principal components were visualised using Manhattan plots.

Pairwise fixation index (FST) values were estimated between all population pairs using R package StAMPP version 1.6.3 (Pembleton *et al*., 2013). Statistical significance was assessed using 100 bootstraps and confidence intervals were calculated at the 95% level. Population structure was further investigated using a model-based clustering approach implemented in ADMIXTURE version 1.90b6.18 (Alexander *et al*., 2009). Input files were prepared from the filtered merged VCF file using PLINK version 1.9 (Chang *et al*., 2015). ADMIXTURE was run for K values ranging from 1 to 10, with ten-fold cross-validation to identify the most likely number of genetic clusters. Each run was performed using four parallel threads. The K value that minimised the cross-validation error was selected as the best-supported number of clusters. Ancestry proportion matrices (Q matrices) were visualised as stacked bar plots using ggplot2, with individuals grouped by sampling region.

Genetic diversity within the collection was estimated as allelic richness using the R package HIERFSTAT version 1.5.11 (Goudet, 2005) with rarefaction to account for unequal sample sizes between regions. Analyses were performed on the merged filtered dataset (1,634 SNPs). As gametophytes are haploid organisms, data were treated accordingly. The genetic diversity available for local breeding programs was assessed by calculating, for each locality the proportion of the total European allelic diversity captured. This was expressed as the number of unique alleles observed across all loci within a given regional subset, divided by the total number of unique alleles observed across the entire collection.

### Genetic crosses and laboratory-based growth of F1 hybrid sporophytes

Individual gametophytes were transferred to PES in 12-well cell culture plates and the filaments were fragmented. Fragments of a single male and a single female gametophyte were then pooled in a 1.5 mL microtube, further fragmented with a micropestle before being transferred to PES in a 55 mm Petri dish, where the mix was cultivated at 10°C under white light from Osram 18W/840 fluorescent “day light” neon tubes (Osram GmbH, Germany) with a 8h:16h light:dark photoperiod to induce fertility. The presence of young F1 sporophytes in the culture was scored after one month. Once the sporophytes were visible to the naked eye, they were transferred to a 140 mm Petri dish and grown to at least 5 mm in length. For each cross, several sporophytes (3 to 15 individuals) were then transferred to a single open-flow low intervention culture system (LICSy) and grown in filtered natural seawater (photoperiod 12h:12h day:night, 11°C) until they had reached a size of 5 to 10 cm and had grown a stipe. Individual sporophytes were then manually attached to 3 mm cotton string and transferred to a 300 L tank with filtered, open-flowing natural seawater and bubbling. Once the sporophytes were 20 to 30 cm long, the cotton strings bearing the sporophytes were transferred to a 4000 L tank with filtered, open-flowing natural seawater and grown up to about 40 cm in length. The light intensity in the 300 L and 4000 L tanks was about 100 µmol photon/m^2^/s (two 5 m rows of white “daylight” 6000 K LEDs and two 5 m rows of white “neutral” 4000 K LEDs).

Photographs were taken weekly to monitor blade surface area in both the LICSy and tank systems but note that individual sporophytes could not be followed in the LICSy system so growth was measured as the total increase in blade surface area for all the individuals in a single culture flask. Blade surface area was calculated using Color Thresholder 1.52a in ImageJ (Schneider *et al*., 2012). For this, the portion of the photograph containing the sporophyte blade was manually delimited and the brightness, contrast, saturation and colour (hue parameter) were automatically adjusted using custom values (determined at the beginning of the experiment based on the first batch of pictures). Blades were detected as particles based on the colour threshold, with the largest particle corresponding to the photographed sporophyte, and their surfaces were measured. Based on these measurements, weekly specific growth rate (SGR) was calculated by comparing blade surface area at the time of measurement (t) with blade surface area one week earlier (t-1) using the formula: 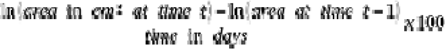 and expressed as the percentage of surface increase per day and cumulative specific growth rate (SGR) was calculated by comparing blade surface area at each time of measurement (t) with blade surface area at the start of the experiment (t0) using the using the formula: 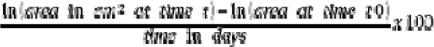 and expressed as the percentage of surface increase per day.

### Estimation of sporophyte growth rates in a common garden free-circulation tank system

F1 hybrid sporophytes from the above 4000 L tank cultures were coppiced by cutting the blade 20 cm above the meristem and all individuals were labelled and cultivated together for two months under a 12h:12h (day:night) light regime with a light intensity of about 100 µmol photon/m^2^/s (two 5 m rows of white “daylight” 6000 K LEDs and two 5 m rows of white “neutral” 4000 K LEDs) in a single 3000 L tank in natural seawater with bubbling and a hydrodynamic current so that the sporophytes circulated freely to avoid bias due to position effects or shading. Blade surface area was measured weekly over the 2-month growing period and weekly SGR was calculated as described in the previous paragraph. At the end of the common garden free-circulation experiment, whole sporophytes were harvested and air-dried by suspending in a large-capacity air drying system (France Haliotis, Plouguerneau, France) for 48 h at 33°C. Each dried sporophyte was then individually reduced to a course-grain powder in a grinder. The ratio of fresh weight (FW) to dry weight (DW) was obtained by weighing the blade after wiping it dry and after drying. The dryness of the sample was estimated on a subsample of about 100 mg by weighing before and after drying for 48h in a ventilated oven (Memmert GmbH, Germany) at 60°C with maximum convection.

### Measurement of alginate and fucoidan content

For each assay replicate, 21 mg of the dried sporophyte material harvested at the end of the common garden free-circulation experiment was transferred to a 2 mL screw-top tube (type E1420-2330; Starlab, Orsay, France) along with three metal beads and ground to a fine powder in a Precellys Evolution (Bertin Technologies, Montigny-le-Bretonneux, France) for four times 20 sec at 5500 rpm. The beads were then removed from the tube and the tubes weighed to obtain the exact weight of powdered material. Two types of analysis were carried out, quantification of neutral monosaccharides and quantification of uronic acids. All biochemical analyses were carried out in triplicate. These were biological replicates whenever possible but for nine crosses biological replicates were not available so technical replicates were carried out (and the type of replicate recorded in each case).

The analyses of monosaccharide composition were carried out using high performance anion exchange chromatography with pulsed amperometric detection (HPAEC-PAD). For the quantification of neutral monosaccharides, samples were hydrolysed with an acid solution in a thermomixer (100°C, 1000 rpm). After appropriate dilution, the resulting hydrolysates were analysed on a CarboPac column under isocratic alkaline conditions. Monosaccharides were identified based on their elution time relative to standard sugar mixes and quantified using response calibration curves for different concentrations of each monosaccharide (Chromeleon software). The same strategy was used for the quantification of uronic acids except that the samples were hydrolysed with an acid solution and then eluted with a sodium acetate gradient.

### Whole genome sequencing and genome-wide variant detection

DNA was extracted from blade tissue of F1 hybrid sporophytes using the Macherey-Nagel Nucleospin Plant II Kit with minor protocol adaptation. Sporophyte tissue was ground in liquid nitrogen using a mortar and pestle. Between 120 and 200 mg of fine powder was then transferred to 3.2 mL of lysis buffer PL1 supplemented with 5% PVP40 (w/v) and prewarmed to 60°C. Lysis was carried out for 10 min at 60°C with 1000 rpm agitation and then at room temperature for 2.5 h. A centrifugation step at 11,000g for 2 min was added before the filtration in order to remove debris and prevent column clogging. An additional wash with PW1 and with PW2 buffer was carried out to further eliminate PCR inhibitors. DNA was eluted into 60 µL of PE buffer at 55°C. DNA concentration and quality was measured using both the NanoDrop One and the Qubit 4 (dsDNA HS kit) and the quality of the extracted DNA was assessed by PCR amplification of an ITS marker using specific primers. The genomic DNA was then sequenced at Novogene (Munich, Germany) on a partial lane of an Illumina NovaSeq X to produce 90 to 120 million 2×150 nt paired end reads per sample. After quality cleaning using Trimmomatic v.0.39 (Bolger *et al*., 2014), the reads were mapped onto the v2 version of the *S. latissima* genome assembly (available at https://phaeoexplorer.sb-roscoff.fr/) using BWA-MEM version 2.2.1 (Li and Durbin, 2009).

To visualise read coverage, normalised coverage ratio was calculated using a two-step process. First the coverage for each 50 kbp window was divided by the mean coverage calculated for all the 50 kbp windows over the entire genome to obtain the normalised coverage. Normalised coverage values were calculated independently for each of the nine F1 hybrid sporophytes and for the reference strain resequencing data using the mean from the corresponding genome in each case. In the second step, the normalised coverage for each 50 kbp window (in terms of chromosome coordinates) along the lengths of each of the nine F1 hybrid sporophyte genomes was divided by the normalised coverage for the corresponding 50 kbp window of the re-sequenced reference genome. This two-step process therefore normalised coverage within each genome and then calculated a coverage ratio compared to the re-sequenced reference genome.

Variants were called independently with DeepVariant version 1.16 (Poplin *et al*., 2018b) and HapoltypeCaller (Poplin *et al*., 2018a) and only variants that were detected by both programs were retained. Variants were quality filtered using bcftools version 1.16 (Danecek *et al*., 2021) based on variant quality (>100) and coverage depth (min 10x, max 100x). When indels were separated by less than 10 bp only one was considered and SNPs within 3bp of an indel were ignored. Only homozygous (defined as frequency >0.9) and heterozygous (defined as 0.4 < frequency <0.6) variants were retained.

To eliminate false positive SNPs and indels, DNA-seq data from the same strain (SLPER63f7) as that used to generate the reference genome assembly was re-mapped onto the reference genome. Analysis of this mapping detected 223,385 SNPs and 160,855 indels (Table S3) that presumably corresponded to sequence errors in the reference assembly. Any variants from the nine F1 hybrid sporophytes that were located at the same coordinates as these spurious variants were removed from the “total variant” datasets to generate clean sets of sequence variants (designated “sequence variants”, Table S3). The clean sequence variants were then used for the subsequent analyses. Note that all counts of variants (SNPs and indels) in this text correspond to counts of numbers of variant loci and do not take into account allelic differences between variants at the same position in the genome.

### Parentage verification

The methodology (in-house scripts) used to verify the parental gametophytes of F1 hybrid sporophytes was developed with assistance from artificial intelligence (Claude opus5 – medium effort – September 2026). Parentage was verified by comparing the genotypes of the F1 hybrid sporophytes with those of their gametophyte parents, together with control non-parental gametophyte strains. Briefly, two sequential tests were applied. First, a single-parental test was carried out to calculate the percentage correspondence of SNPs between the genotype of each sporophyte and that of each gametophyte strain. This first analysis was restricted to sites that were homozygous in the relevant sporophyte genome to allow one-for-one comparisons at each site. Second, a biparental test was carried out that calculated the percentage of sporophyte SNPs, both heterozygous and homozygous, that were coherent with the expected genotype of a combined pair of gametophyte strains. To limit this analysis to a manageable number of gametophyte pairs, it was carried out using random samples of four gametophytes from each of the 33 populations (with the exception of Point de Cabellou, where only one gametophyte strain was available), including the expected parental strains of the nine F1 hybrid sporophytes when ddRAD-seq genotype data was available (i.e. for 13 of the 18 parental gametophytes). The variant data generated by Stacks based on the gametophyte ddRAD-seq data set the reference allele (REF) as the major allele in the analysed population. To compare with the F1 hybrid sporophyte data, for which REF had been determined relative to the reference genome assembly, it was therefore necessary to adjust the reference allele for the gametophyte data so that the reference allele corresponded to the allele present in the reference genome assembly. Two filtering steps were carried out to identify informative sites. These filters were applied independently for each comparison. The first filter selected only ddRAD-based gametophyte variants that had also been called in the relevant sporophyte. The second filter was different for the two tests described above. For the first test the filter selected only sites that were homozygous in the relevant sporophyte. For the second test, the filter only selected alleles that did not correspond to the dominant allele in the gametophytic population (i.e. selection of alleles with a within-population frequency <0.5). This second filter was intended to decrease noise by reducing chance matches due to the prevalence of the major alleles.

## RESULTS

### Establishment, sexing and genotyping of a collection of European *S. latissima* gametophyte strains

Wild *S. latissima* sporophytes were sampled across 33 sites in eight countries, spanning the European range of this species, during two successive sampling campaigns carried out in the context of the European projects Genialg and SeaMark (Fig. 1, Table S4). Sporulation of the sporophytes allowed the established a large collection of 650 gametophyte strains (444 for Genialg and 206 for SeaMark; Table S4). A PCR-based approach was used to sex the gametophyte strains and 583 of the strains were genotyped (444 for Genialg and 139 for SeaMark; Table S4) using the ddRAD-seq method (Mauger and Avia, 2024). The ddRAD-seq genotyping was carried out in two separate experiments (ddRAD pool 1, 409 gametophyte strains, and ddRAD pool 2, 174 gametophyte strains, see Materials and Methods). The two ddRAD-seq experiments used different restriction enzyme combination, so they were therefore initially analysed independently and the datasets were then merged to identify and analyse the set of common variant loci. After removal of indels, filtering for mean read depth (5×–40×), minimum allele count of 2, and retention only of biallelic sites, 141,735 and 214,915 SNPs were retained for the ddRAD pool 1 and ddRAD pool 2 datasets, respectively. The two filtered datasets were then merged based on shared genomic coordinates and additional filtering was applied for missing data per individual (>5%), missing data per locus (>5%), heterozygosity per individual (>5%), heterozygosity per locus (>5%) and to retain only one SNP per RAD-tag. The final merged dataset consisted of 1,634 SNPs genotyped across 366 individuals.

### Genetic structure of the gametophyte collection

Principal component analysis of the 366 individuals based on the 1,634 SNPs of the merged dataset identified two distinct clusters, a southern cluster with populations from Spain, Portugal, France and southern England and a northern cluster grouping populations from Norway, Sweden, Ireland, Scotland and the Faroe Islands (Fig. 1B). Within the northern group, the Faroe Islands populations were located closer in the PCA to populations in Scotland and Ireland than to populations in Norway and Sweden. When the contribution of individual SNPs to the first and second principal components was examined values were broadly and uniformly distributed across the genome, with no evidence of localised peaks on specific chromosomal regions (Fig. S1). This pattern is consistent with differentiation driven primarily by genetic drift and isolation, rather than by divergent selection, which would be expected to result in elevated values at discrete genomic loci.

ADMIXTURE analysis of the 1,634 SNPs in the merged dataset identified K=4 as the most likely number of genetic clusters (Fig. 1C), corresponding to an Iberian cluster (Spain, Portugal), an Atlantic and English Channel cluster (Brittany, northern France, southern England), a northern Atlantic cluster (Ireland, Scotland, Faroe Islands), and a Scandinavian cluster (Norway, Sweden). Individual ancestry assignments were highly consistent within each cluster, with most individuals showing more than 95% membership to a single cluster and minimal admixture between groups (Fig. 1C). A small number of Norwegian individuals showed minor ancestry contributions from the northern Atlantic cluster, consistent with the geographic proximity of these regions.

Pairwise F_ST_ values were estimated between all population pairs using the 1,634 SNPs in the merged dataset. The vast majority of pairwise F_ST_ values were statistically significant (p < 0.05, 100 bootstraps), indicating genetic differentiation among populations across the European range of *S. latissima*. F_ST_ values were consistent with the four-cluster structure identified by the ADMIXTURE analysis (Fig. S2). Within-group differentiation was moderate, with mean pairwise F_ST_ values of 0.20 for Scandinavian populations (Norway and Sweden), 0.24 for northern Atlantic populations (Ireland, Scotland and Faroe Islands), 0.18 for Atlantic populations (UK/France/Brittany) and 0.29 for Iberian populations (Spain and Portugal). Between-group differentiation was substantially higher, with mean F_ST_ values of 0.34 between the Scandinavian and northern Atlantic clusters, 0.41 between the Scandinavian and Atlantic clusters and 0.34 between the northern Atlantic and Atlantic clusters. The highest differentiation was observed between the Iberian cluster and all other populations (mean F_ST_ 0.55), confirming the marked genetic distinctiveness of southern European populations.

Allelic richness varied among sampling regions (Table S5), with the highest values observed in southern Brittany (1.36 allele per locus, rarefaction corrected), followed by the Faroe Islands (1.21) and Norway (1.10). Combining local populations increased this proportion significantly: pooling of three Breton populations (LEC, PEN, STG) captured 64.5% of total diversity, while combining Faroe Islands and Scottish populations (FUN2, GTA2, FWT) or combining Norwegian populations (NAV, BAR, SOT) captured 58.2% and 48.6%, respectively. These results indicate that pooling of germplasm from populations within the same genetic cluster would allow approximately half of the total allelic diversity to be retained compared with the diversity available across the entire European range of *S. latissima*.

### Production of F1 hybrid sporophytes by crossing gametophyte strains

Crosses were carried out between defined pairs of male and female gametophytes to evaluate the phenotypic characteristics of F1 hybrid sporophytes produced by different combinations of parental strains. To compare the characteristics of F1 hybrid sporophytes derived from narrow and wide genetic crosses, these experiments focused on two geographical regions, the southern part of the Finistère region of Brittany, France and the Faroe Islands. Gametophyte strains from these two regions were crossed in both intra- and inter-population combinations (Fig. 2A). Altogether, four classes of cross were carried out: intra-population crosses in which either both parents were from Brittany (designated ALO crosses) or both parents were from the Faroe Islands (designated ORF) and inter-population crosses between Breton and Faroe Islands gametophyte strains, in which either the Breton (designated ALF) or the Faroe Islands (designated ORA) strain was the female parent (Fig. 2A). A total of 168 crosses was carried out and 143 of these crosses (85.1%) produced sporophyte offspring (Fig. 2B, Table S6). The gametophyte strains collected as part of the Genialg project had been maintained in a vegetative state for up to ten years following isolation (Table S6). These strains were therefore able to recover fertility even after this prolonged storage under red light conditions.

**Fig. 2.**
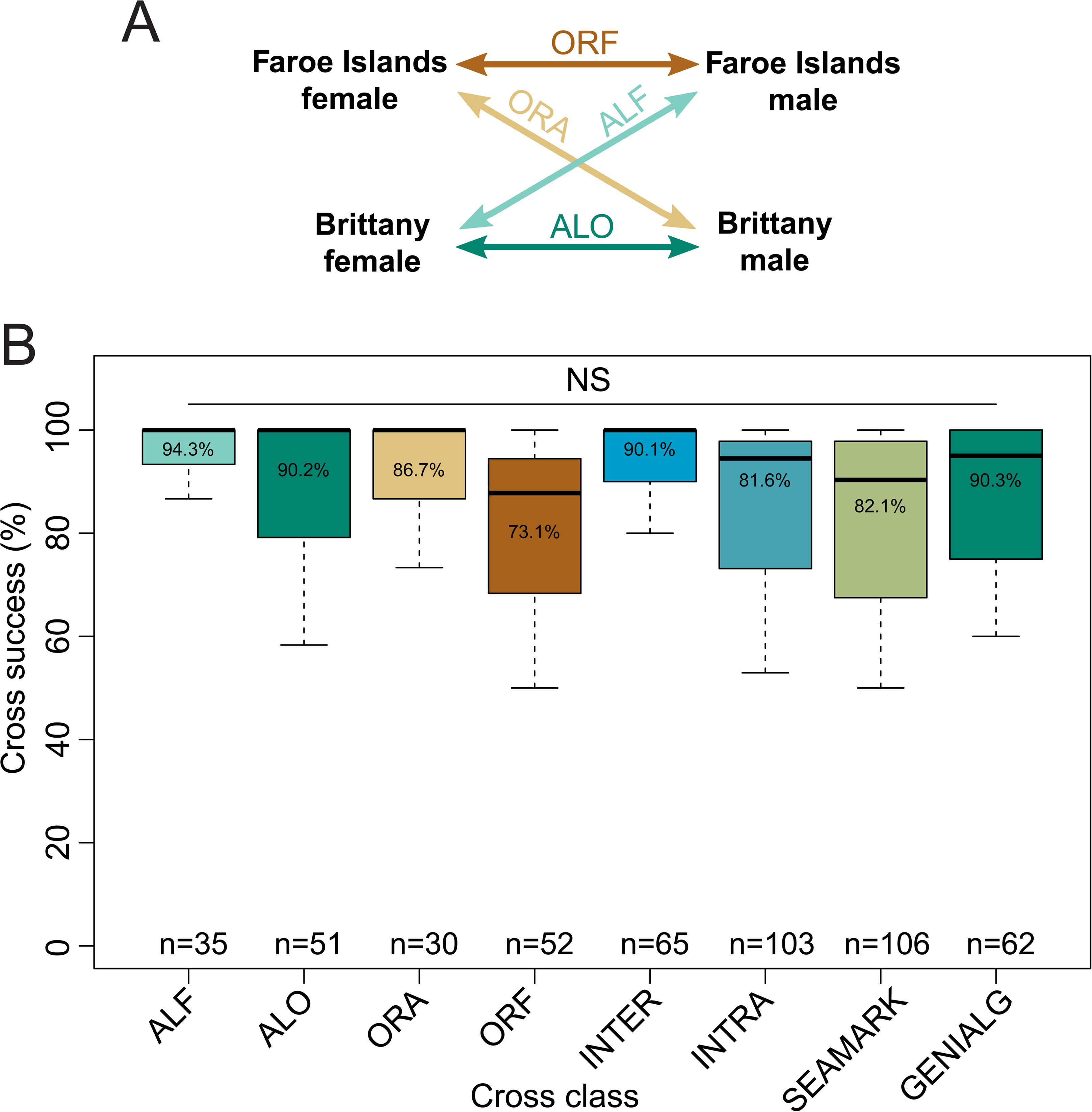
Crossing scheme and cross success. A. Diagram showing the four classes of cross that were carried out. ALO and ORF were intra-population crosses, ALF and ORA were inter-population crosses. B. Percentages of different types of cross that successfully produced sporophytes. SEAMARK and GENIALG indicate the strains isolated during the two European projects. Note that the Genialg strains have been maintained in culture for up to ten years. n, number of crosses; NS, no significant difference; ALO, Brittany x Brittany cross; ORF, Faroe Islands x Faroe Islands cross; ALF, Brittany x Faroe Islands cross; ORA, Faroe Islands x Brittany cross, INTER, inter-population cross; INTRA, intra-population cross.

### F1 hybrid sporophytes exhibited marked differences in growth performance

Sixty of the crosses that produced sporophyte progeny were further cultured and analysed. The F1 hybrid sporophytes were initially raised in Petri dishes, then transferred as pools of sporophytes from an individual cross to a low-intervention culture system (LICSy) and then grown as individual sporophytes attached to lines in 300 L and 4000 L tank systems (Table S6). Measurements of blade surface area made during cultivation in the LICSy and in the 300 L and 4000 L tank systems indicated marked differences in the growth rates of different hybrid F1 sporophytes (Fig. S3, Table S6). However, these analyses only provided approximate comparisons because it was necessary to grow the sporophytes in several batches and, therefore, it was not possible to ensure that all individuals were grown under exactly the same conditions. For example, different individuals would have experienced different levels of shading, especially during the later stages of growth and it is also possible that there were differences in the exact composition of the natural seawater during the course of the experiment. To address these problems, the growth performance of the F1 hybrid sporophytes was tested in a single common garden experiment. For this, 116 sporophytes, derived for 50 different crosses (18 ALO, 11 ORF, 10 ALF crosses and 11 ORA), were coppiced by cutting the blade 20 cm above the meristem and all the individuals were then cultivated together in a 3000 L tank. The individuals were labelled but not attached, allowing them to circulate freely in the tank in the air-bubbled seawater medium, avoiding position or shading bias. Blade surface area was monitored weekly over a 2-month period. An almost four-fold difference in growth rate (SGR) was observed between the least and the best performing F1 hybrid sporophytes (Fig. 3A).

**Fig. 3.**
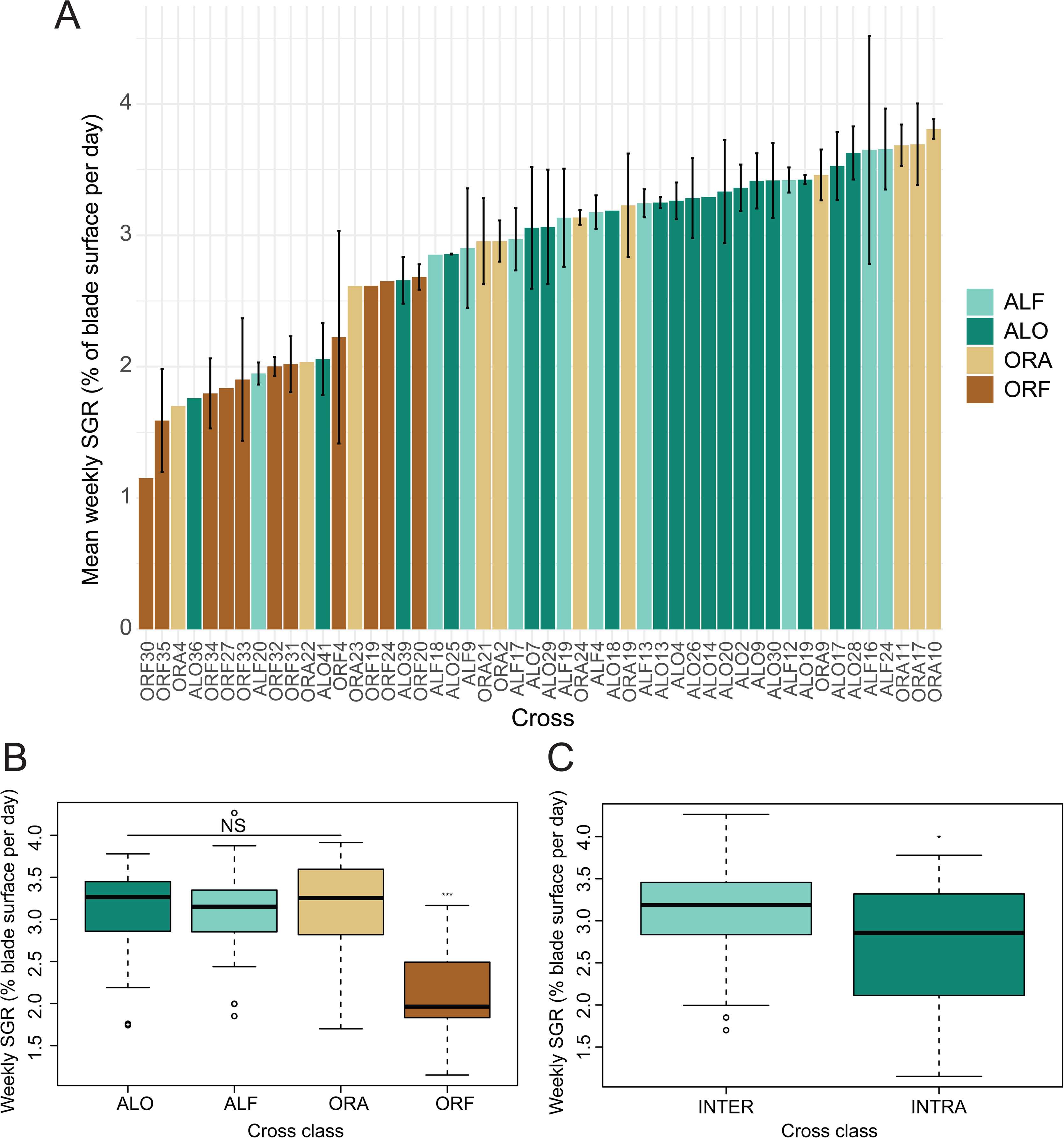
Growth performance of F1 hybrid sporophytes in a common garden tank growth experiment. A. Mean daily growth rates (increase in blade surface) for F1 sporophytes derived from different types of crosses grown in the common garden experiment. Mean daily increases in blade surface were calculated for individual sporophytes and the average of the mean daily growth rates calculated over the growth period was plotted, together with the variance (standard error). B. Distributions and medians of daily growth rates for the four different classes of cross. C. Distributions and medians of daily growth rates for intra- and inter-population crosses. ALF, Brittany x Faroe Islands; ALO, Brittany x Brittany; ORA, Faroe Islands x Brittany; ORF, Faroe Islands x Faroe Islands; INTER, inter-population; INTRA, intra-population; NS, not significant; *, *p*-value ≤ 0.05; ***, *p*-value ≤ 0.01.

Sporophytes derived from crosses between two parents that were both from the Faroe Islands (cross class ORF) performed less well, in terms of SGR, than any of the other classes of cross indicating that the culture conditions used for this experiment may not have been optimal for strains from the Faroe Islands. For example, F1 sporophytes were cultivated under a 12h:12h (day:night) photoperiod whereas natural day length in the Faroe Islands varies between 5h and 20h year-round, with very short days in the winter. Importantly, however, no significant differences were detected between inter- or intra-population crosses involving one or two parental strains from Brittany, respectively (Fig. 3B). Note also that variance in weekly SGR did not appear to be markedly greater for inter-population crosses involving Breton strains compared to Breton intra-population crosses (Fig. 3B) indicating that inter-population crosses do not appear to be more likely to produce high-performance outliers than intra-population crosses (Fig. 3C). Overall, therefore, this experiment indicated that inter-population crosses did not produce significantly faster growing strains than intra-population crosses.

### Cell wall polysaccharide composition

Quantifications of neutral monosaccharides and of uronic acids were carried out on F1 hybrid sporophyte material derived from 24 crosses (six ALO, six ORF, six ALF and six ORA) following cultivation in the common garden free-circulation tank experiment. Faroe Islands intra-population hybrid sporophytes (ORF crosses) contained significantly higher levels of neutral monosaccharides than Breton intra-population hybrid sporophytes (ALO crosses) (Fig. 4A, Table S7). Inter-population crosses between strains from the two regions (ALF and ORA) had levels that were intermediate between those of the two classes of intra-population cross (ALO and ORF). Analyses of individual neutral monosaccharides indicated that the ORF hybrids accumulated significantly higher levels of mannitol and fucose (Fig. 4B-D, Table S7). Xylose accumulated to significantly lower levels in intra-population crosses between Breton strains (ALO) than for any of the other classes of cross (Fig. 4E, Table S7).

**Fig. 4.**
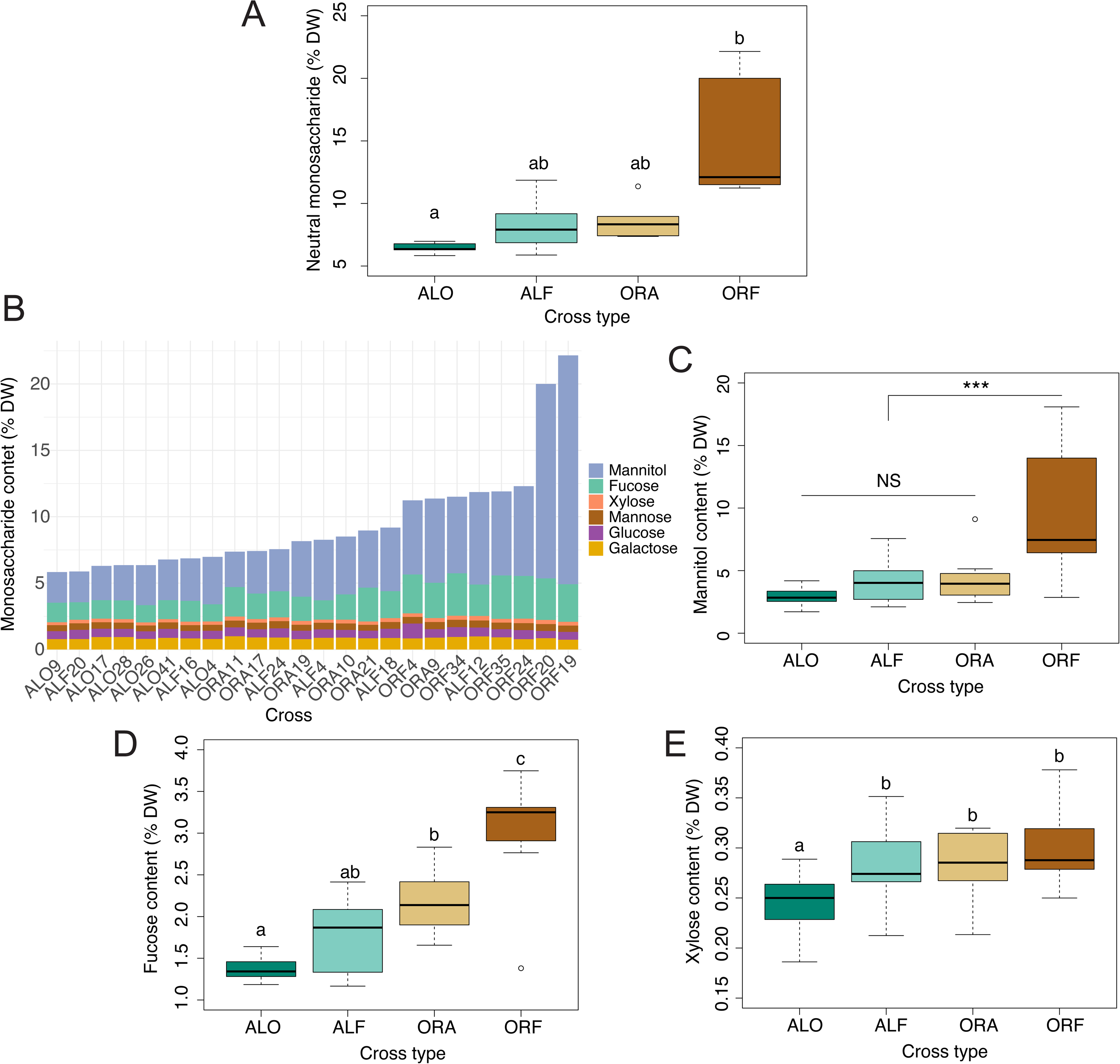
Neutral monosaccharide content of F1 hybrid sporophytes in a common garden tank experiment. A. Distribution of neutral monosaccharide content (as a percentage of dry weight) across the four classes of genetic cross. B. Neutral monosaccharide content of F1 hybrids as a percentage of dry weight. C. Distribution of mannitol content across the four classes of cross. D. Distribution of the fucose content of the F1 hybrids grown in the common garden experiment plotted as a percentage of dry weight. E. Distribution of the xylose content of the F1 hybrids grown in the common garden experiment plotted as a percentage of dry weight. Differences between crosses were analysed using the non-parametric Kruskal-wallis test followed by pairwise comparison using a Dunn test. Letters denote significant differences (*p*-value < 0.001 Bonferroni corrected for A. and *p*-value < 0.05 Bonferroni corrected for B. to E.). DW, dry weight; NS, not significant; ***, *p*-value ≤ 0.01; ALF, Brittany x Faroe Islands; ALO, Brittany x Brittany; ORA, Faroe Islands x Brittany; ORF, Faroe Islands x Faroe Islands; INTER, inter-population; INTRA, intra-population.

Alginic acid content varied about 0.3-fold across the assayed samples (Fig. 5A, Table S7), with inter-population Brittany x Faroe Islands crosses (ALF) having significantly higher levels (Fig. 5B, Table S7) than intra-population Faroe Islands x Faroe Islands crosses (ORF). Similarly, the ratio of guluronic acid to mannuronic acid varied across samples (Fig. 5C, Table S7) and, again, inter-population Brittany x Faroe Islands crosses (ALF) had significantly higher ratios (Fig. 5D, Table S7) than intra-population Faroe Islands x Faroe Islands crosses (ORF).

**Fig. 5.**
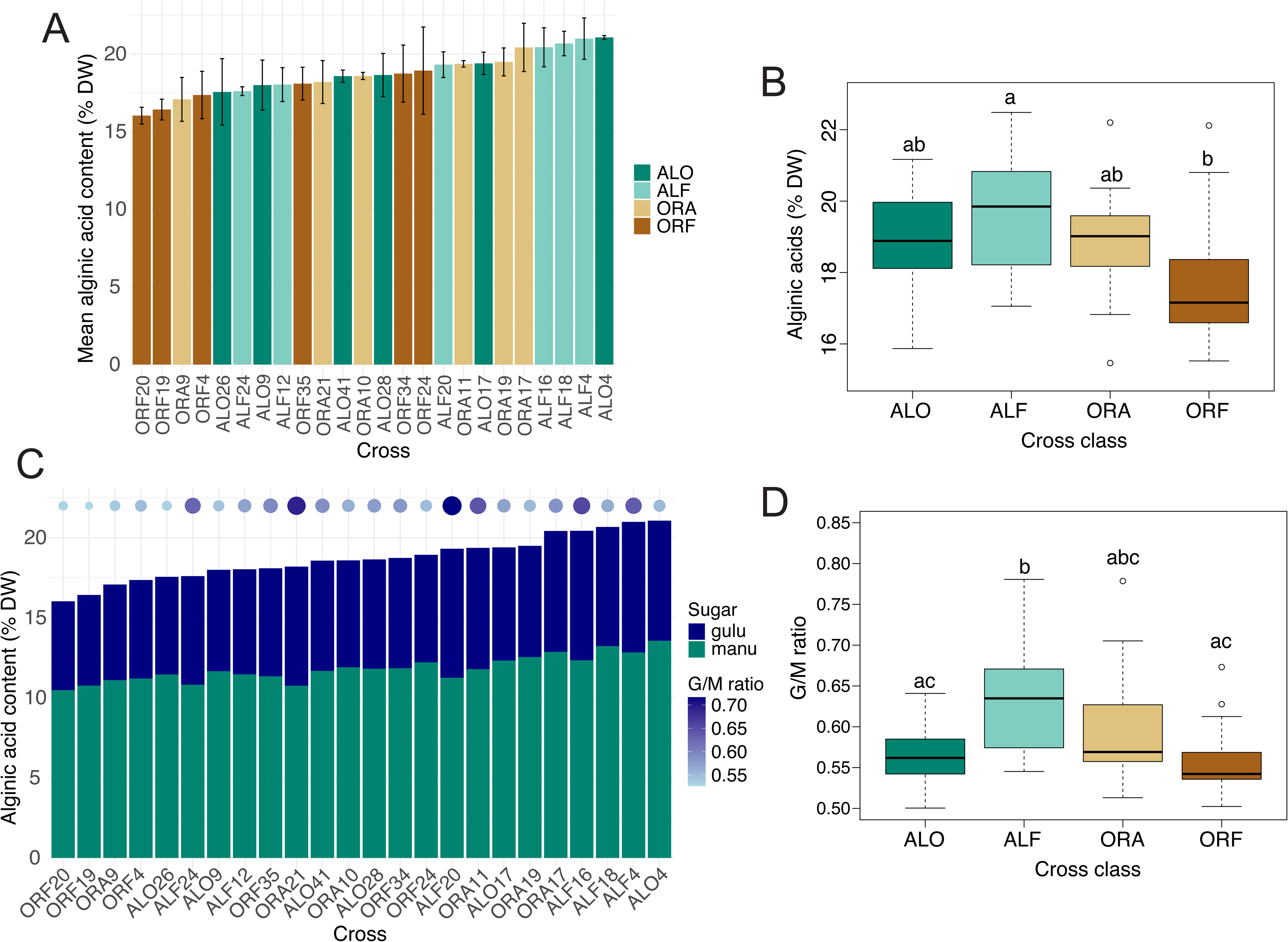
Alginic acid content and composition in F1 hybrid sporophytes in a common garden tank experiment. A. Alginic acid content as a percentage of dry weight. B. Alginic acid content across the four classes of genetic cross. C. Composition of the alginic acid of the F1 hybrid sporophytes grown in the common garden experiment. Blue discs indicate the ratio of guluronic acid to mannuronic acid. D. Distributions of guluronic acid to mannuronic acid ratios across the four classes of genetic cross. Letters denote significant differences (*p*-value < 0.05 Bonferroni corrected). DW, dry weight; ALF, Brittany x Faroe Islands; ALO, Brittany x Brittany; ORA, Faroe Islands x Brittany; ORF, Faroe Islands x Faroe Islands; gulu, guluronic acid; manu, mannuronic acid; G/M, ratio of guluronic acid to mannuronic acid.

Overall, therefore, there were not marked differences in neutral monosaccharide and uronic acids content across the crosses but some significant differences were detected, related to the geographic origins of the parental strains. However, as in the growth performance tests above, there was no evidence for marked differences in neutral monosaccharide and uronic acids content between sporophytes derived from wide crosses and sporophytes derived from intra-population crosses. The observation that there were some significant differences that were related to geographic origin indicates that the differences had a genetic basis, which could be exploited in future breeding programs. However, it is possible that at least part of the difference between neural monosaccharides and uronic acids contents for Faroe Islands x Faroe Islands crosses (ORF) compared with the other classes of cross may have been due to the lower growth rates of sporophytes derived from this class of cross under the experimental conditions used here (Fig. 3B).

### Genome sequencing of selected, rapidly growing F1 hybrid sporophytes

Nine of the F1 hybrid sporophyte strains that had shown rapid growth rates were selected for more complete genetic characterisation using a whole genome sequencing (WGS) approach. The selected strains included five sporophytes derived from Brittany x Brittany crosses (ALO) and four sporophytes derived from Faroe Islands x Faroe Islands crosses (ORF; Table S8). Mapping of the Illumina short read WGS data to the *S. latissima* v2 genome assembly (Table S9) did not detect any large-scale insertions or deletions across the strains, apart from an apparent duplication of a region at the end of chromosome 12 in strain ALO20 (Fig. S4). Variant calling identified between 0.80 and 1.63 million SNPs and 0.22 and 0.38 million indels per F1 hybrid sporophyte strain (Table S3). The majority of the SNPs detected were specific to either the Breton or the Faroe Islands populations (Fig. S5). Greater numbers of variants were detected in strains derived from ORF crosses than in strains derived from ALO crosses (Table S3). This difference was expected because the Faroe Island (ORF) strains are more distantly related to the reference genome, than are the Breton (ALO) strains (Table S8).

The WGS data was also used to assess whether ddRAD genotyping captures a representative subset of genome-wide sequence variants. In general, this does seem to be the case. The spectra of SNP mutations were similar for the two methodologies, with transitions being about twice as abundant as transversions (Fig. S6). Moreover, both the WGS- and ddRAD-derived SNPs were distributed evenly along the *S. latissima* chromosomes, although at a lower density for the ddRAD-derived SNPs, as expected (Fig. S7). Both WGS- and ddRAD-derived SNPs were enriched in intergenic regions and depleted from exons and coding regions (Fig. S8, Tables S10 and S11), although ddRAD SNPs were less depleted from coding regions (Fig. S8C).

The high levels of heterozygosity detected for some of the gametophyte strains suggested that a proportion of the isolated gametophytes may not have been clonal. This observation prompted us to carry out parentage verifications for the F1 hybrid sporophytes by comparing sporophyte genotype data with the genotypes of the parental gametophytes (Fig. S9). ddRAD-seq-based genotype data was available for the gametophyte parents of all of the F1 hybrid sporophytes, except for ORF30 and ORF35, where neither parent had been genotyped, and ALO36, where only the female parent had been genotyped. For the seven F1 hybrid sporophytes that could be analysed, the expected parents were confirmed in five cases (although only the female parent could be verified for ALO36). Of the remaining two F1 hybrid sporophytes, ALO20 appeared to be a doubled haploid with the female parent (GA_STG_57_F10) genotype (or possibly a self cross with a highly similar contaminating brother of GA_STG_57_F10) and it was only for ORF4 that both of the expected parents were rejected as progenitors. Note that, although genotype data was not available for the parents of ORF30 and ORF35, the analysis confirmed the population of origin of these F1 hybrid sporophytes. To summarise, therefore, this analysis only provided clear evidence of misattributed paternity/maternity for one of the seven analysable crosses, indicating that the observed heterozygosity did not markedly impact the outcome of the crossing experiment.

## DISCUSSION

This study has established a large collection of *S. latissima* gametophyte strains derived from 33 sites along the coastline of western Europe and has generated ddRAD genotype data for 90% of the collection. The results of the analysis of genetic structure across this collection were consistent with several previous studies in that they identified a clear genetic differentiation between populations from northern and southern Europe (Luttikhuizen *et al*., 2018; Guzinski *et al*., 2020; Diehl *et al*., 2023; Jaugeon *et al*., 2025). The genetic structure analysis indicated that the populations in this study could be grouped into four distinct clusters (Fig. 1C), likely reflecting divergent phylogeographic histories, with Iberian strains being the most genetically differentiated. Substructure was also apparent within each cluster, with substantial differentiation among strains (F_ST_ values of up to 0.2). Further sampling and studies will be required to determine whether some of this within-cluster variation reflects local adaptation among strains and may be relevant to future breeding programs.

Analysis of the ddRAD-seq data identified significant levels of heterozygosity in some of the gametophyte isolates, particularly from the SeaMark sampling campaign. This heterozygosity suggests that many of the isolated strains are not pure clones and, therefore, that it would be advisable to carry out an additional gametophyte purification step when using these strains for future experimentation. In general, care should be taken to verify parental genotypes when analysing the progeny of crosses, even when working with isolated gametophyte strains.

Crossing experiments were carried out using strains from two distant geographic localities. These experiments showed that fertility could be efficiently induced in strains that had been stored for more than ten years under red light conditions. This observation confirms previous studies, which have shown that kelp gametophytes can retain fertility after prolonged periods of cultivation (tom Dieck (Bartsch) and de Oliveira, 1993; Bartsch, 2018; Martins *et al*., 2019; Sæther *et al*., 2024), in one case even after 45 years in culture (*S. latissima* culture 3301/3302 isolated by Klaus Lüning in 1974 from Oslofjord and employed in crosses by Pearson *et al*., 2019). Maintenance of gametophytes in vegetative culture is therefore an effective and secure means of biobanking genetic resources for future breeding programs.

Comparisons of growth rates of F1 hybrid sporophytes derived from intra- and inter-population crosses between the strains from the two localities - the Finistère region of Brittany and the Faroe Islands - did not indicate any marked advantage of wide crosses compared to crosses between strains from the same locality. This is also an interesting result because it indicates that restricting breeding programs to the use of locally-sources strains is not expected to limit the effectiveness of the breeding process. Similarly, comparisons of neutral monosaccharide and uronic acids content of individuals derived for intra- and inter-population crosses did not indicate that the wide crosses produced hybrids with more extreme or improved characteristics.

Taken together, therefore, the analyses carried out in this study indicate that breeding strategies focused uniquely on the use of locally-sources genotypes would not necessarily be less efficient that a classic approach involving wide crosses. This is an important conclusion because breeding is a promising approach to optimise multiple aspects of kelp farming, including biomass yield, content of specific valuable compounds and biomolecules, susceptibility to biofouling, disease resistance and suitability for specific aquaculture procedures such as harvesting or mechanical coppicing. The results reported here indicate that it should be possible to work towards these objectives without resorting to wide crosses and therefore to the introduction of foreign genotypes into cultivation sites. Note, however, that inbreeding depression has been reported for *S. latissima* (Bråtelund *et al*., 2026a) so care should be taken to ensure that sufficient genetic diversity is present in the locally-sourced germplasm used for breeding and that selfing be avoided if possible. In this context, the robust observation from genetic analyses that European *S. latissima* populations fall into two principal clusters, corresponding to northern and southern populations (Neiva *et al*., 2018; Guzinski *et al*., 2020; Diehl *et al*., 2023; Jaugeon *et al*., 2025; Mauger *et al*., 2026 and this study), indicates that breeding programs could possibly integrate germplasm sources from quite broad geographic areas (either northern or southern Europe) and still avoid the introduction of foreign genotypes.

Interestingly, Cohen *et al*. (2025), who carried out wide crosses between *S. latissima* strains, did not observe a correlation between heterosis (in terms of morphological traits) and geographic distance, supporting the idea that crossing genetically distant strains may not be more effective as a breeding approach to increase biomass. However, they indicated that geographic distance may not have been necessarily correlated with genetic distance in their study.

The F1 hybrid sporophytes generated by this study exhibited a broad range of growth rates, indicating that the genetic composition of the parent gametophytes markedly influenced the growth capacity of the sporophyte progeny. Comparable ranges of growth rates were observed for sets of individuals derived from intra- or from inter-population crosses. It was therefore possible to select strains with high growth rates issued from intra-population crosses that will be suitable for open-sea deployment under mariculture conditions at the two study sites. The high-growth-rate strains generated in this study were derived from pairs of parental gametophytes that can be multiplied vegetatively. providing the possibility of generating large populations of identical, rapidly growing F1 hybrid sporophytes for cultivation. Genome-wide sequence variant catalogues were generated for nine of the strains with high growth rates, providing precise genetic descriptions of these improved strains. This comprehensive genetic information is expected to be useful for the future deployment of genetics-based approaches to breed improved strains of *S. latissima* for mariculture.

## CRediT authorship contribution statement

**Zofia Nehr**: Conceptualization, Investigation, Data curation, Resources, Formal analysis, Visualization, Writing – original draft, Writing – review and editing. **Francis Le Madec**: Investigation, Writing – review and editing. **Olivier Godfroy**: Investigation, Formal analysis, Data curation, Visualization, Writing – review and editing. **Aurélie Préchoux**: Investigation, Writing – review and editing. **Alan Le Moan**: Formal analysis, Writing – review and editing. **Komlan Avia**: Formal analysis, Writing – review and editing. **Dan Potin**: Investigation, Writing – review and editing. **Martina Strittmatter**: Investigation, Writing – review and editing. **Marie-Mathilde Perrineau**: Resources. **Cecilia Rad-Menendez** Resources, Writing – review and editing. **Delphine Scornet:** Resources, Writing – review and editing. **Bertrand Jacquemin**: Resources, Writing – review and editing. **Emilie Gouhier**: Resources, **Jerome Coudret**: Resources, Writing – review and editing. **Lucie Jaugeon**: Resources, Writing – review and editing. **Stephane Mauger**: Resources, Writing – review and editing. **Carla Ruiz Gonzalez**: Resources, Writing – review and editing. **Myriam Valero**: Supervision, Writing – review and editing. **Claire Gachon**: Supervision, Writing – review and editing. **Philippe Potin**: Conceptualization, Project administration, Supervision, Writing – review and editing. **J. Mark Cock**: Conceptualization, Project administration, Supervision, Data curation, Writing – original draft, Writing – review and editing.

## Data availability

The sequence data used in this study have been deposited in the European Bioinformatics Institute/European Nucleotide Archive (EBI/ENA) database under the project accessions PRJEB86118 and PRJEB86958 for the gametophyte ddRAD-seq data (this study), PRJEB111452 for the F1 hybrid sporophyte genome resequencing data (this study) and ERR12682613 for the *S. latissima* gametophyte strain SLPER63f7 DNA-seq data (Denoeud *et al*., 2024). See Tables S4 and S8 for details. VCF files for both the gametophyte ddRAD-seq genotyping and the F1 hybrid sporophyte whole genome sequencing have been deposited in the CNRS Research Data dataset “Replication Data for: Comparison of the performance of sporophytes derived from narrow and wide genetic crosses between strains of *Saccharina latissima*” (https://doi.org/10.57745/JDTNQ1). The v2 version of the *S. latissima* genome assembly is available at https://phaeoexplorer.sb-roscoff.fr/.

## Acknowledgments

We thank Sunniva Tangen Haldorsen, Floor Marsman, Inga Maria Mikkelsen, Ólavur Gregersen, Mathilde Lemoine, Laurent Lévèque, Wilfried Thomas, Yann Fontana, Mathieu Oriot, Isabel Azevedo, Phil Kerrison, Lars Brunner, Wouter Visch, Akira Peters, Antoine Fort and Thea Helen Johnsplass for help with the sporophyte sampling, Claire Daguin, Marion Ballenghien and the Roscoff Genomer platform for help with ddRAD library preparation, the Roscoff Aquarium Service for access to kelp culture infrastructure, the UMR8227 Culture Service for help with strain maintenance, Sylvain Huchette at France Haliotis for access to their seaweed drying chamber and the Culture Collection of Algae and Protozoa (in particular Callum O’Connell) and the Roscoff Culture Collection for strain biobanking. We are grateful to the Roscoff Bioinformatics platform ABiMS (http://abims.sb-roscoff.fr), which is part of the Institut Français de Bioinformatique (ANR-11-INBS-0013) and BioGenouest network, for providing both help and computing and storage resources.

## Supplementary figure legends

**Fig. S1**. Manhattan plot of absolute loading illustrating the contribution 1,634 SNPs to principal components 1 (top panel) and 2 (bottom panel) of the PCA shown in Fig. 1B. The X-axes indicate the position of each SNP along the 31 chromosomes of the *S. latissima* genome.

**Fig. S2**. Pairwise F_ST_ matrix. White lines delineate the four clusters of populations identified by the ADMIXTURE analysis. ***, *p*-value < 0.05.

**Fig. S3**. Growth performance of F1 hybrid sporophytes in the LICSy system and in the 300 L and 4000 L tank systems. A. Average weekly SGR of the F1 sporophytes for each cross in the LICSy system. B. Comparison of weekly SGRs for the four classes of cross in the LICSy system. C. Comparison of weekly SGRs for inter- and intra-population crosses in the LICSy system. D. Average weekly specific growth rate of the F1 sporophytes for each cross in the 300 L and 4000 L tank systems. E. Comparison of weekly SGRs for the four classes of cross in the 300 L and 4000 L tank systems. F. Comparison of weekly SGRs for inter- and intra-population crosses in the 300 L and 4000 L tank systems. n, number of crosses; NS, no significant difference; **, *p*-value < 0.005; ALO, Brittany x Brittany cross; ORF, Faroe Islands x Faroe Islands cross; ALF, Brittany x Faroe Islands cross; ORA, Faroe Islands x Brittany cross, INTER, inter-population cross; INTRA, intra-population cross.

**Fig. S4**. WGS DNA-seq data coverage analysis. Normalised coverage ratios over 50 kbp windows. Each individual dot corresponds to a 50 kbp window and the line is a mobile mean covering 200 kbp (four 50 kbp windows). A. Overview showing normalised coverage ratios along the 31 chromosomes of the *S. latissima* genome for each of the nine strains. B. Detailed view of the SL_12 chromosome showing a region of higher read coverage at the end of the chromosome for the strain ALO20.

**Fig. S5**. Overlaps of SNP datasets detected by WGS and ddRAD-seq. A. Venn diagram showing the overlap of SNPs detected by either in ddRAD (gray) or WGS in at least one ALO strain (green) and at least one ORF strain (brown). B. and C. Upset diagrams showing the overlap of SNPs identified in the four ORF strains (B) and the five ALO strains (C).

**Fig. S6**. Characteristics of sequence variants detected by WGS. Histograms show either the numbers of SNPs corresponding to each type of nucleotide change (upper panels) or the number of indels corresponding to each length of insertion (positive) or deletion (negative) (lower panels). Data is shown for each of the nine F1 hybrid sporophyte WGS analyses and for the ddRAD data. Indels were not analysed for the ddRAD data.

**Fig. S7**. Distribution along the genome of SNPs detected using either WGS or ddRAD-seq. SNP density per kilobase was calculated in 10 kbp windows (individual dots) along the 31 chromosomes of the *S. latissima* genome for both the SNPS detected using ddRAD (grey) and the SNPs detected in at least one of the nine strains using WGS (blue). The plotted line is a tendency curve based on a Generalized Additive Model (“gam” method of the “geom_sooth” tool from ggplot2 R package), the dashed line indicates mobile means over 200 kbp (20 point) intervals.

**Fig. S8**. Localisation of WGS-detected variants. A. and B. Localisation of the identified SNPs in the exonic, intronic and intergenic regions of the genome, reported as either absolute numbers (A) or percentages (B). C. Proportions and numbers of SNPs present in coding regions (CDSs). The dashed line marks the proportion of the genomic sequence corresponding to CDS (0.0463).

**Fig. S9**. Genotype-based verification of the parental gametophytes of F1 hybrid sporophytes. A. Genetic similarity scores for comparisons between F1 hybrid sporophytes and individual gametophyte strains. Numbers indicate the percentage of informative sporophyte homozygous sites at which the gametophyte carries the same allele. A site was considered to be informative only when the sporophyte allele was a minor allele within the gametophyte’s population. B. Identification of the best-fitting gametophyte parent pairs based on ranking of the Mendelian compatibility of every gametophyte pair from the panel tested (741 mother x father pairs per sporophyte). The top eight pairs are shown. Note that genotype data was not available for the parent gametophytes of ORF30 (GA_GTA2_94 and GA_FUN2_97) and ORF35 (GA_FUN2_101 and GA_GTA2_7) nor for the male parent of ALO36 (GA_LEC_20). Gametophyte names have been shortened, e.g. LEC_5 corresponds to strain GA_LEC_5.

## Supplementary tables

**Table S1**. PCR primers used to sex gametophyte strains.

**Table S2**. Mean percent heterozygosity for each gametophyte strain.

**Table S3.** Sequence variants detected using WGS and ddRAD sequencing.

**Table S4**. List of isolated gametophyte strains and accession numbers for ddRAD-seq data.

**Table S5**. Allelic richness of populations and regions.

**Table S6**. List of the crosses carried out between gametophyte strains and growth characteristics of crosses in the LICSy system, in 300 L and 4000 L tank systems and in the common garden free-circulation 3000 L tank experiment.

**Table S7.** HPAEC-PAD quantification of neutral monosaccharides and uronic acids.

**Table S8.** Accession numbers for WGS DNA-seq sequencing data.

**Table S9.** Mapping of WGS DNA-seq sequence data to the reference (version v2) *S. latissima* genome.

**Table S10.** Distribution of WGS SNPs across genome features.

**Table S11.** Sequence variants are underrepresented in coding region and exons.

