## Supplementary figures and images for "Comparison of the performance of sporophytes derived from narrow and wide genetic crosses between strains of the cultivated kelp *Saccharina latissima*"

### Supplementary figure 1

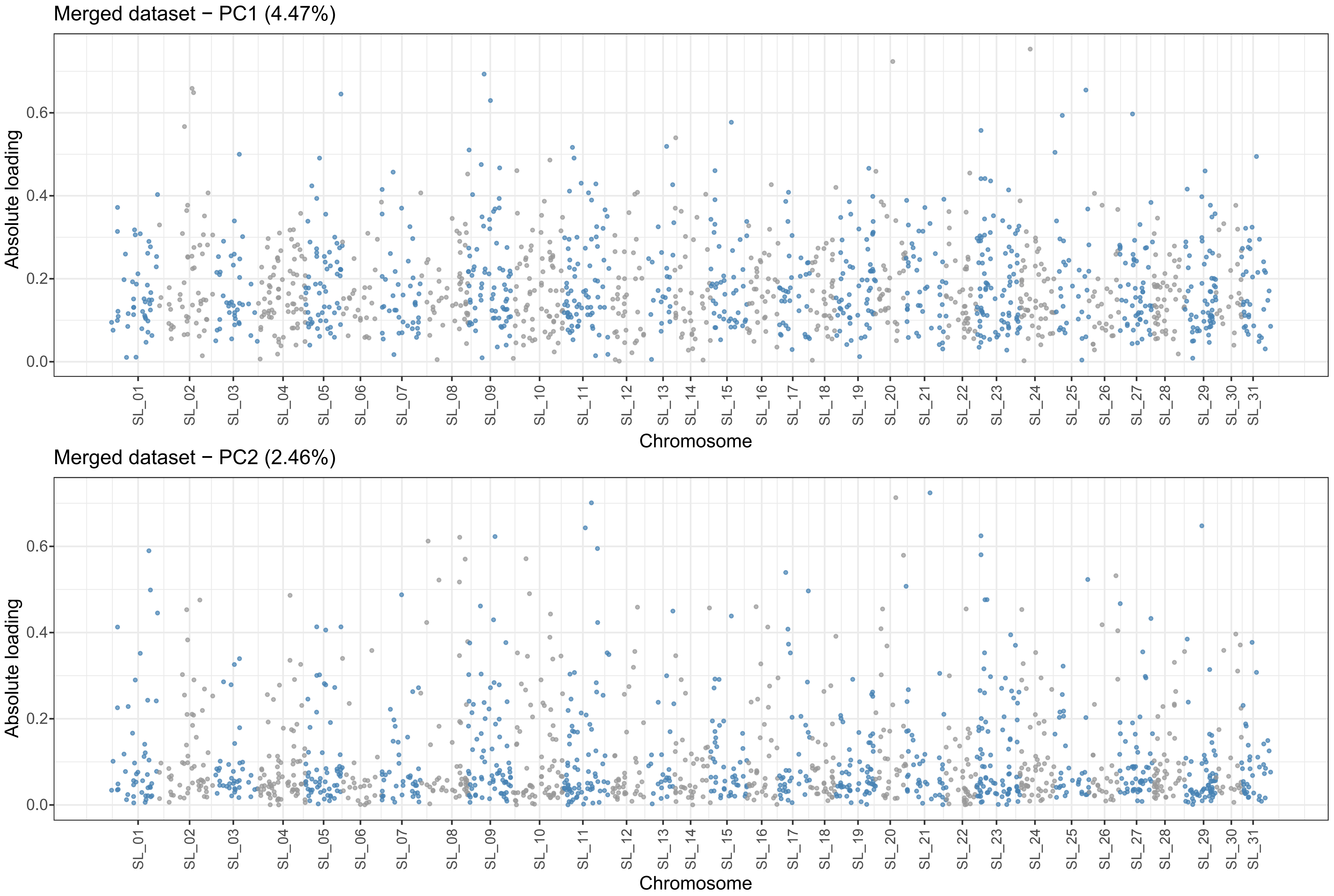

### Supplementary figure 2

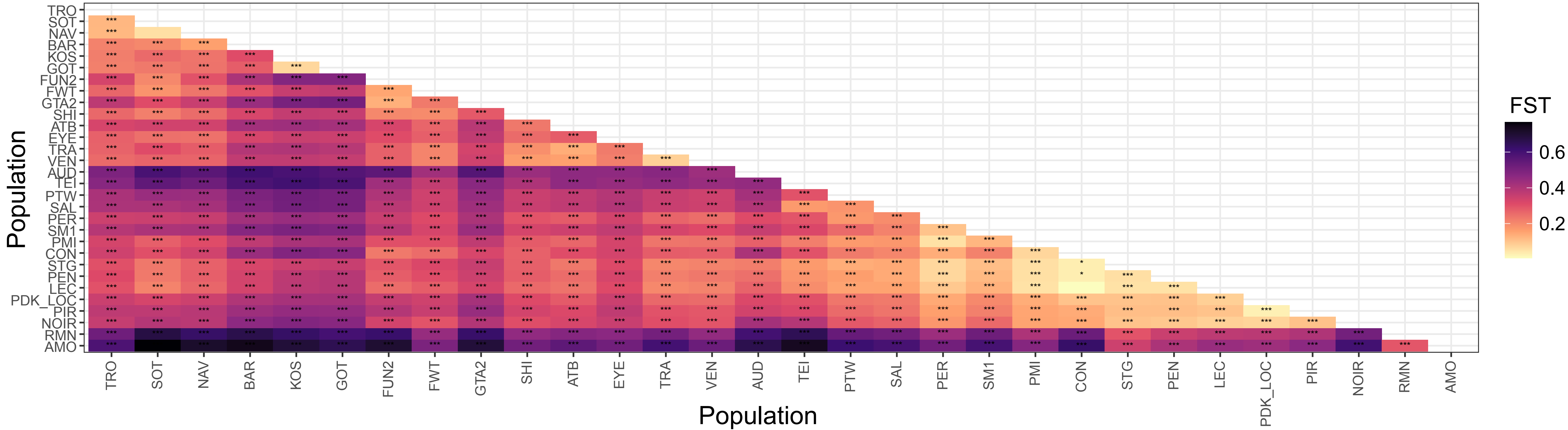

### Supplementary figure 3

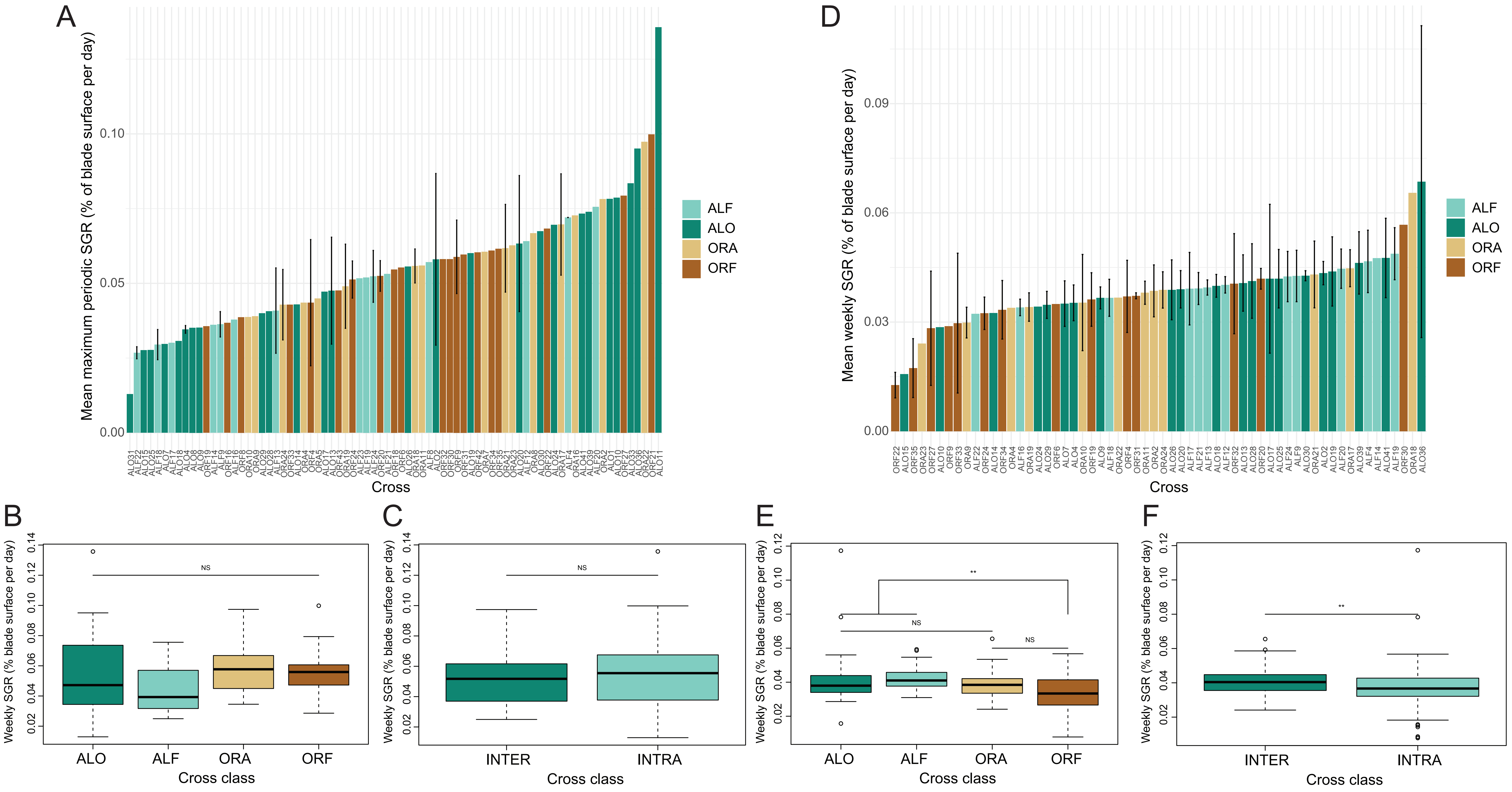

### Supplementary figure 4

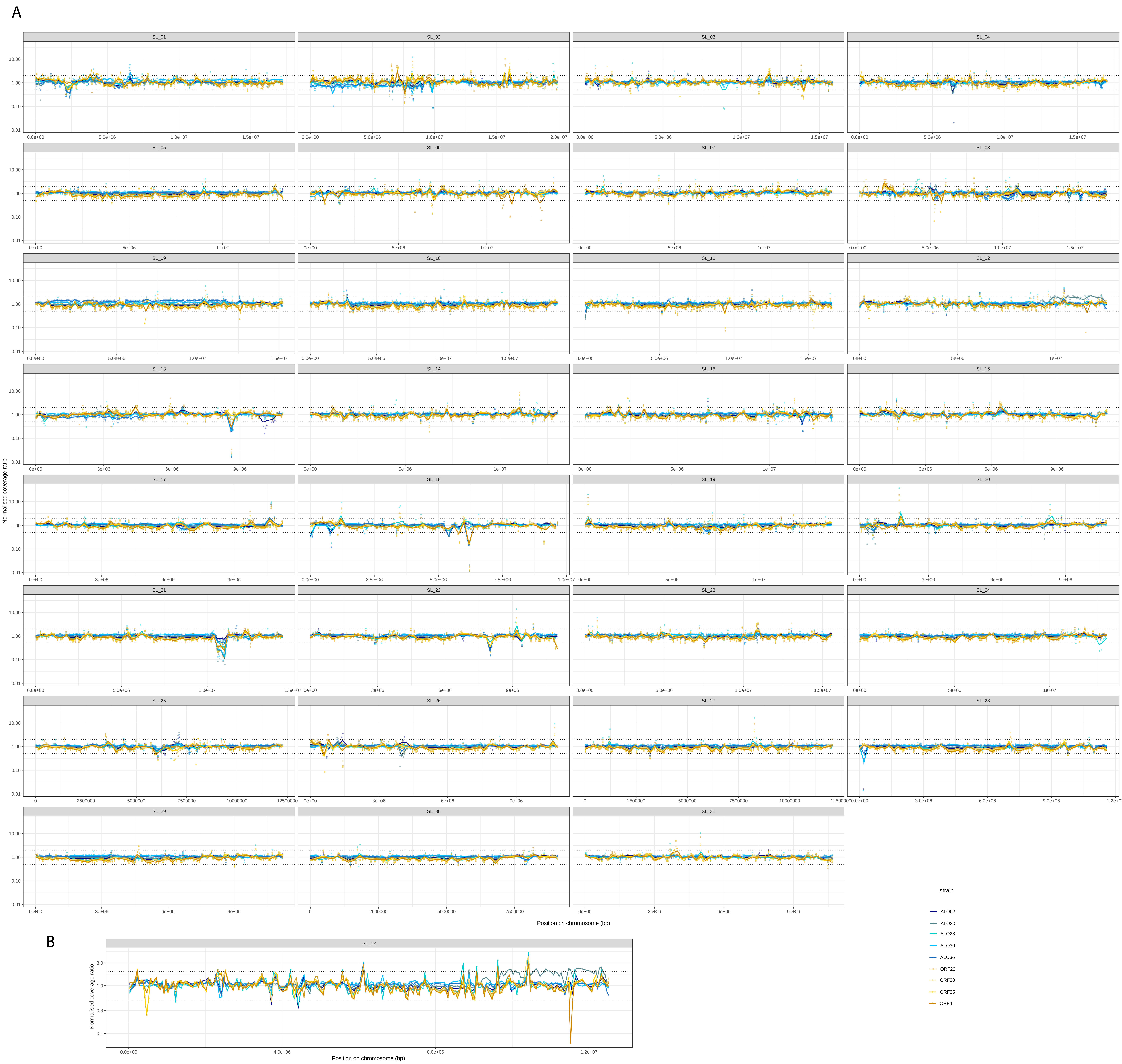

### Supplementary figure 5

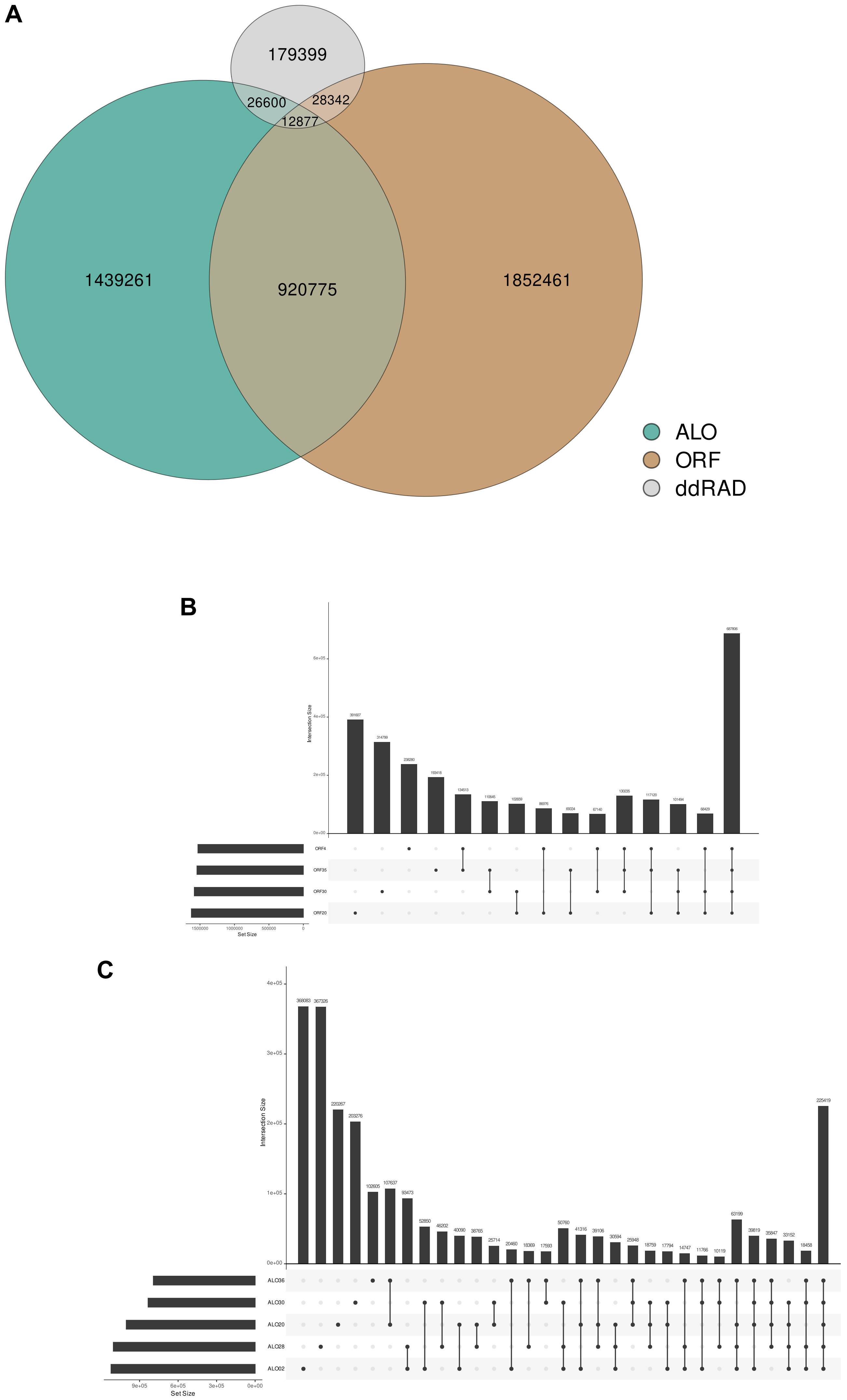

### Supplementary figure 6

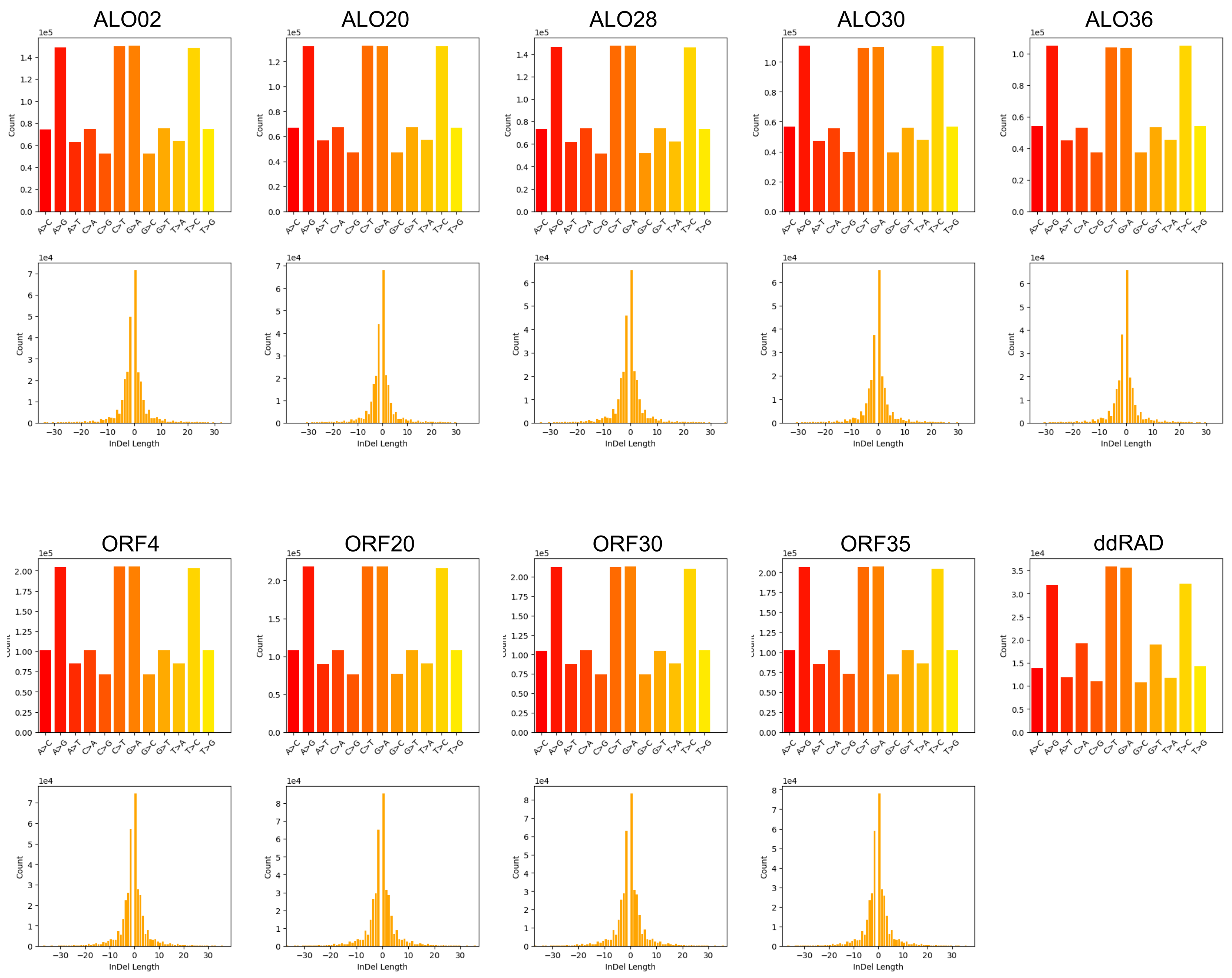

### Supplementary figure 7

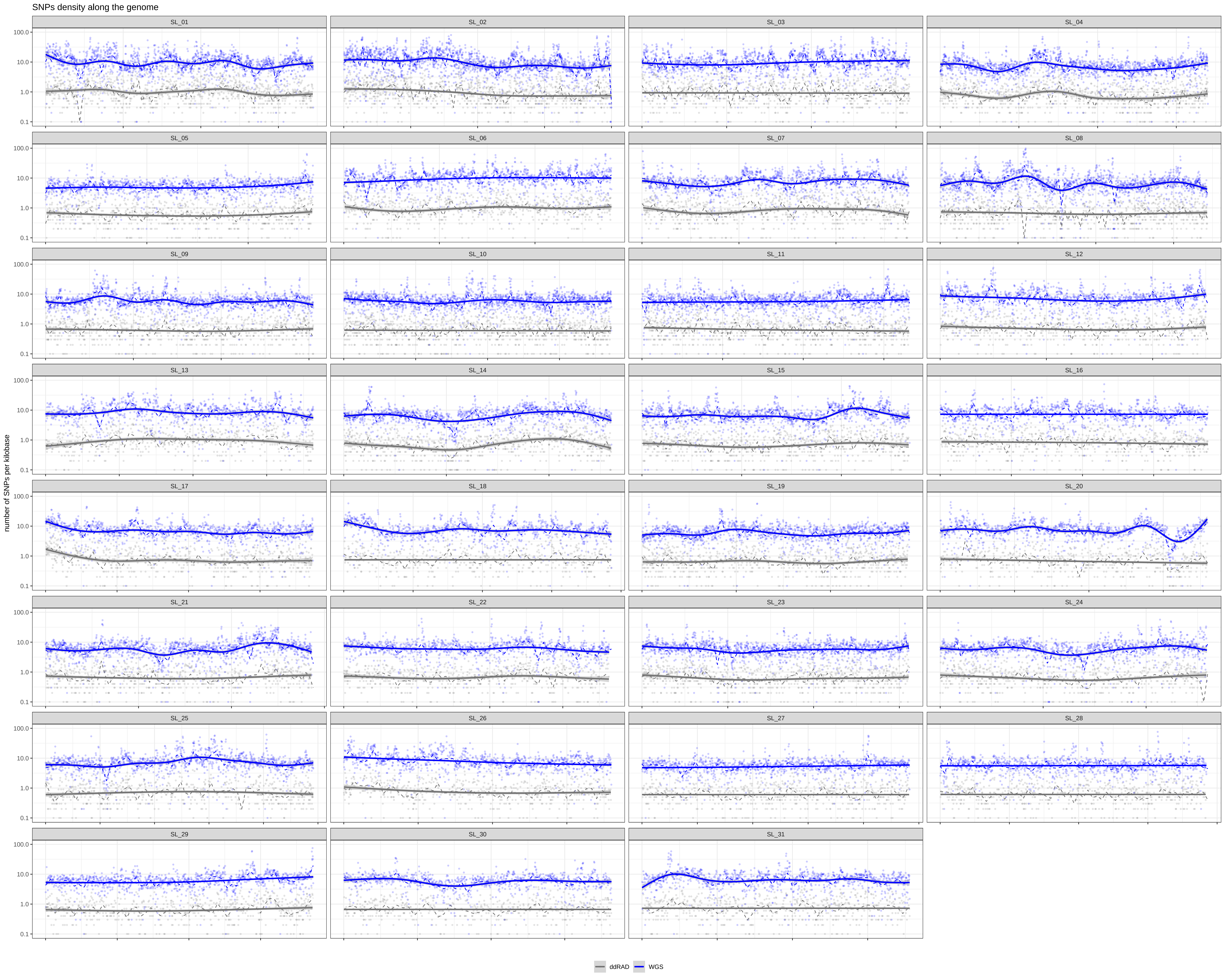

### Supplementary figure 8

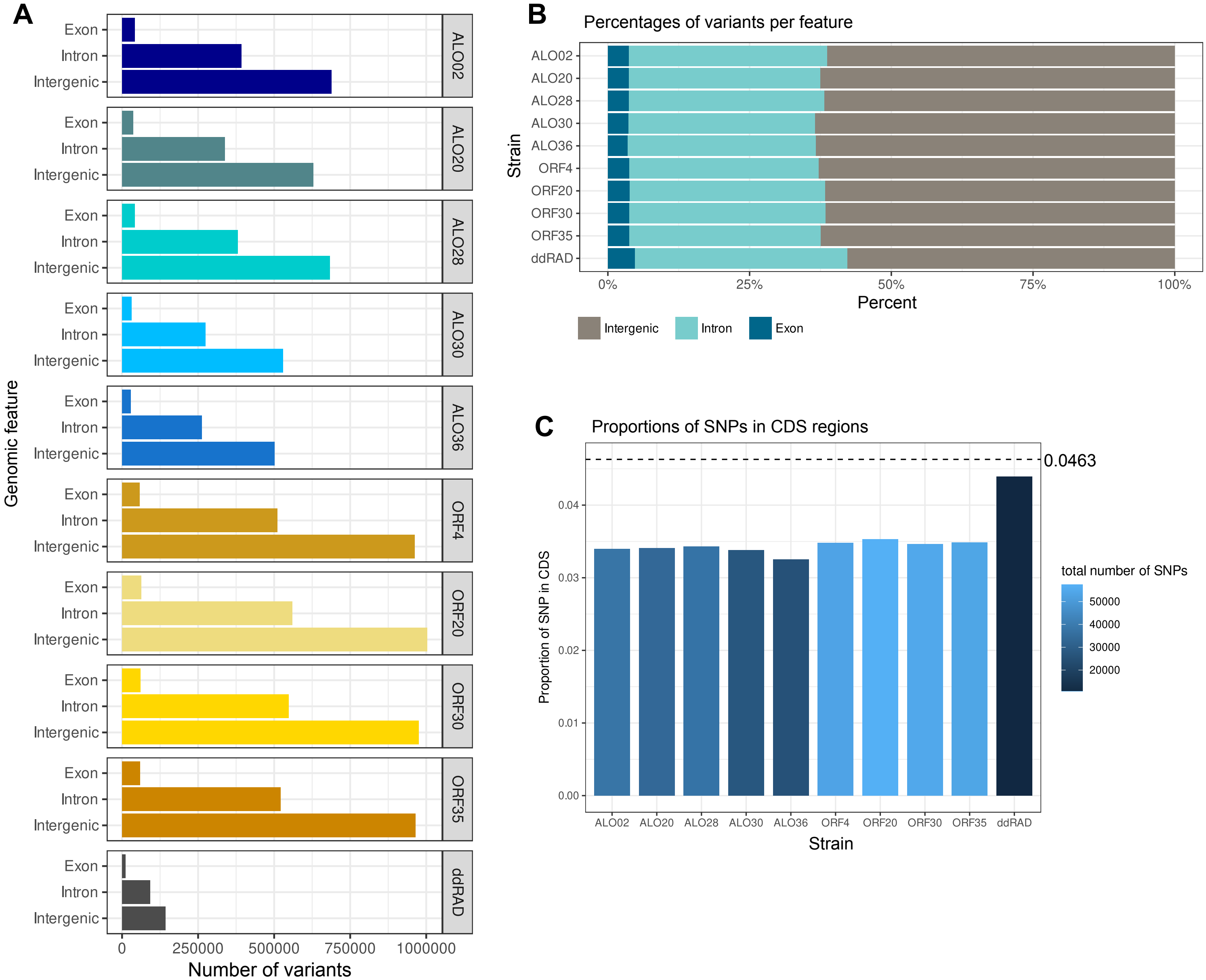

### Supplementary figure 9

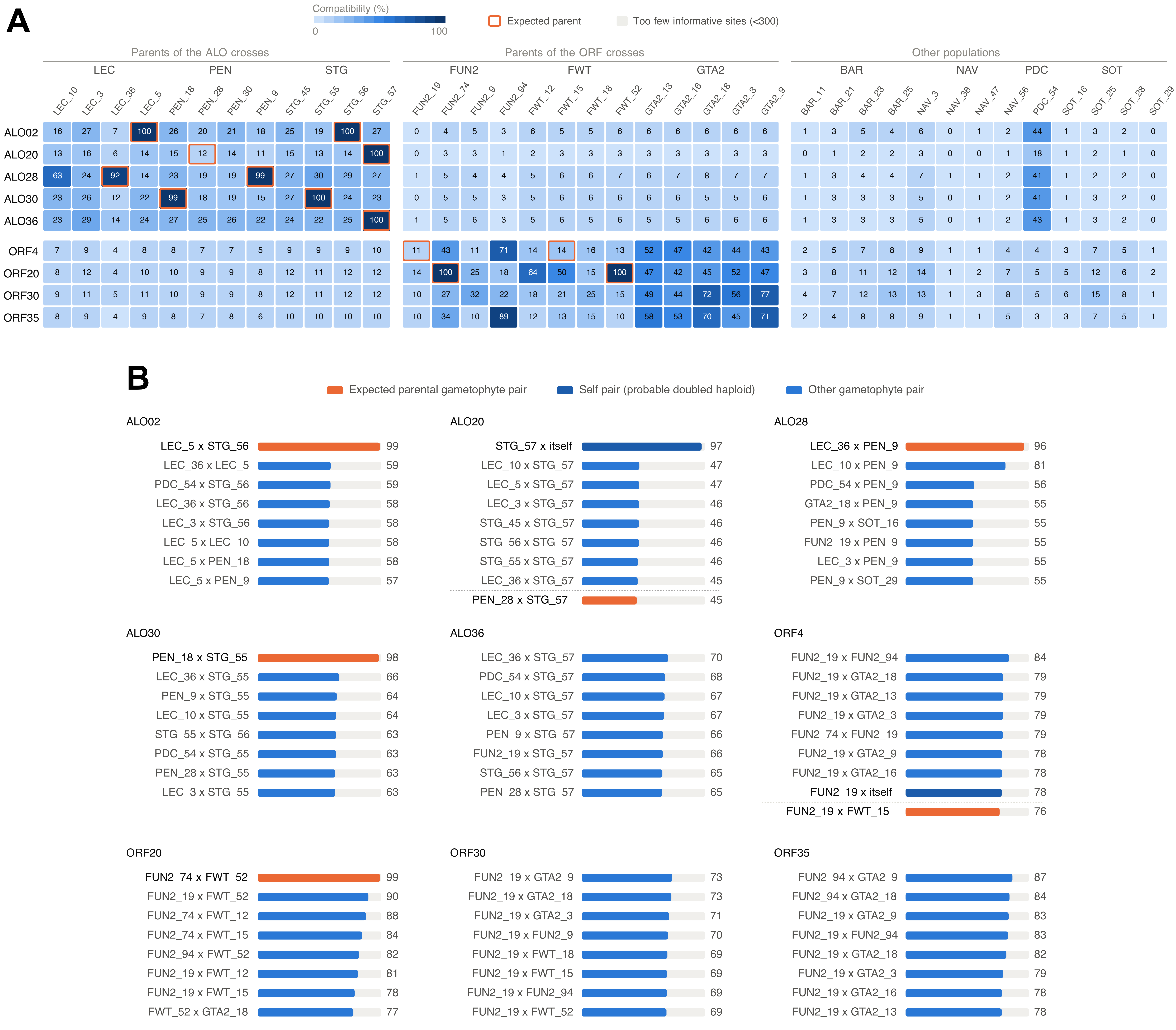
